# Early Prediction of Preeclampsia Based on Transposable Elements signature in cell-free RNA

**DOI:** 10.1101/2024.11.08.622691

**Authors:** Manthan Patel, Ahmed S. Ali, Adrianna Dabrowska, Fanny Boulet, Ajay Kumar Sinha, Madapura M Pradeepa

## Abstract

Preeclampsia (PE) is a pregnancy-associated hypertension disorder affecting 5–10% of pregnant women each year worldwide, which leads to adverse maternal and child outcomes. PE remains inadequately predicted, prevented and treated. Here, we comprehensively investigated altered transcriptomic and epigenetic changes in the PE placenta and maternal cell-free RNA (cfRNA). We show that many transposable elements (TEs) in sub-families are deregulated in the gestational age-matched PE placenta. Increased expression of endogenous retrovirus (ERV) and long interspersed nuclear element 1 (L1, LINE1) subfamilies of TEs is associated with higher histone acetylation levels at their regulatory elements. A higher TE transcript level correlates with the type I interferon (IFN-I) pathway, suggesting inflammation associated with PE could be due to the sensing of TE transcripts by the antiviral innate immune system. Consistent with higher TE transcript levels in the PE placentas, analysis of maternal cfRNA data revealed differential levels of TE subfamilies in PE compared to healthy controls. Machine learning-based training for TE transcripts for early prediction of PE showed a robust performance in the validation cohort, with an area under the curve (AUC) of 0.88, 81% sensitivity, and 74% positive predictive value (PPV). Overall, we show TE deregulation in the placenta is associated with PE, and maternal cfRNA TE signature accurately predicts early diagnosis of PE, which can improve prophylaxis and obstetric outcomes.

## Introduction

Preeclampsia (PE) is a serious and complex pregnancy-specific hypertensive disorder affecting mothers and infants. PE accounts for 14% of maternal deaths each year and is a major cause of premature births ^1,2^. PE can also lead to other complications, including HELLP syndrome, premature birth, foetal growth restriction and stillbirth. The human placenta has a compressed lifespan, governs pregnancy outcomes, and plays a vital role in the offspring’s health. Abnormal placentation leads to pregnancy complications, including PE, early onset (<34 weeks) PE (EOPE), PE with intrauterine growth restriction (PE+IUGR) or severe PE (∼2% of the pregnancies). Although mechanisms of pathophysiology are unclear, epigenetic changes in the placenta are linked to PE ^3,4^. Early prediction of subsequent development of PE before 16-week gestation in pregnancy is hugely vital as prophylaxis at this stage prevents preterm PE-associated complications ^5,6^.

The loss of heterochromatin and DNA methylation associated with ageing, senescence, cancer and neurological diseases causes deregulation of L1, HERV and *Alu* TE families, which leads to elevated IFN-I innate immune pathway ^7–15^. Similarly, in the Drosophila ageing model, stimulating retrotransposon activity increases mortality and accelerates a subset of ageing phenotypes ^16^. The cytotrophoblast epigenome is shown to be dramatically reprogrammed during pregnancy, which is associated with an increased global level of DNA methylation and heterochromatic marks (H3K9me3) with gestational age (GA). In contrast, the H3K27ac level reduces with GA ^4^. H4K16ac is a highly abundant mark enriched at euchromatin, particularly at gene bodies, enhancers and TEs ^17,18^. H4K16ac is implicated in ageing and senescence, as its level increases at specific genomic loci with age and senescent mammalian cells ^19–21^. We have recently demonstrated that H4K16ac regulates the transcription of TEs in embryonic stem cells ^18^. TEs are repressed in somatic tissues but are detected at higher levels during early development, embryonic stem cells and placenta ^22–24^. The preeclamptic placenta displays senescence and accelerated ageing phenotype, together with associated DNA methylation and gene expression signatures ^3,25,26^. Interestingly, recent findings show the role of lysine acetyltransferases p300 and KAT8 in trophoblast stem cell proliferation and differentiation and invasion properties of TSCs ^27–29^. Acetyl-CoA metabolism maintains histone acetylation levels, including H4K16ac, essential for differentiating STBs from TSCs ^27^. Further investigations on the mechanism through histone acetylation axis regulate trophoblast lineage in the placenta development and placental disorders.

A better understanding of the transcriptomic and epigenetic changes associated with PE will aid in early predicting pregnancy complications. Recent proof-of-concept works have demonstrated that maternal circulating cell-free RNA (cfRNA) and cell-free DNA (cfDNA) sequencing data can identify PE-associated transcripts and DNA methylation signatures that predict PE ^30–33^.

Transposable elements (TEs) constitute nearly 53% of the human genome and have significantly contributed to rewiring the gene-regulatory landscape by acting as species- and tissue-specific transcriptional enhancers ^18,34–39^. Long terminal repeats (LTRs) of endogenous ERVs are exapted to function as tissue-specific enhancers, including placental-specific enhancers ^18,34,40–43^. Moreover, ERV envelope-derived genes such as Syncytins have co-opted the fusogenic role in villous cytotrophoblasts (VCTs) to form multinucleated syncytiotrophoblasts (STBs) ^44^. Despite these intriguing connections between TEs and placental development, evolution and function, epigenetic factors contributing to TE regulation in the placenta and the effect of TE deregulation are unclear.

We discovered that TEs are specifically deregulated in PE and associated with heightened IFN-I pathway activity, indicating that TE deregulation may play a role in the development of PE. Analysing maternal cfRNAseq datasets showed that TEs exhibit PE-specific differential levels in maternal circulation. Additionally, through a machine learning approach, we identified TE transcript signatures that can predict PE in the early stages of pregnancy.

## Results

We aimed to investigate altered transcriptomics and epigenetic landscape of PE placenta; we recruited preterm/term preeclampsia (PE) or PE with intrauterine growth restriction (PE+IUGR) and gestational age-matched and unmatched healthy normotensive controls (Extended Data Table 1). To ensure the observed findings are not due to differences in the gestational age of the placenta, we used two independent RNAseq cohorts that are gestational age-matched (Figs. 1a and 1b, and Extended Data Fig. 1a,b) ^45–48^. The foetal side of the placental biopsies containing chorionic villus tissue was banked to investigate altered histone acetylation and transcriptomic changes associated with PE. We performed polyA-enriched RNAseq for two independent areas of the placenta per biopsy. We analysed RNAseq data from our cohort separately and by integrating with public datasets that are gestational age-matched placenta (GSE114691, 21 control, 20 PE and 20 PE+IUGR) and GSE186257, 18 control and 26 severe-PE). Differential expression analysis showed expected upregulation of PE-specific transcripts such as INHBA, ENG, FSTL3, LEPTIN, HTRA4 and FLT1 in PE placentas from public and our cohort (Fig. 1c, and 1d, Extended Data Fig. 1). The findings were consistent when comparing gene expression changes in PE for two independent cohorts with GA matched controls (Extended Data Fig. 1a-d). Gene ontology analysis for the DEGs revealed enrichment of angiogenesis, hypoxia, inflammatory response, EVT cell types and metabolic pathways, including glycolysis, which are known to contribute to the altered Acetyl-CoA levels (Extended Data Fig. 1e and 1f) ^49^.

**Fig. 1:**
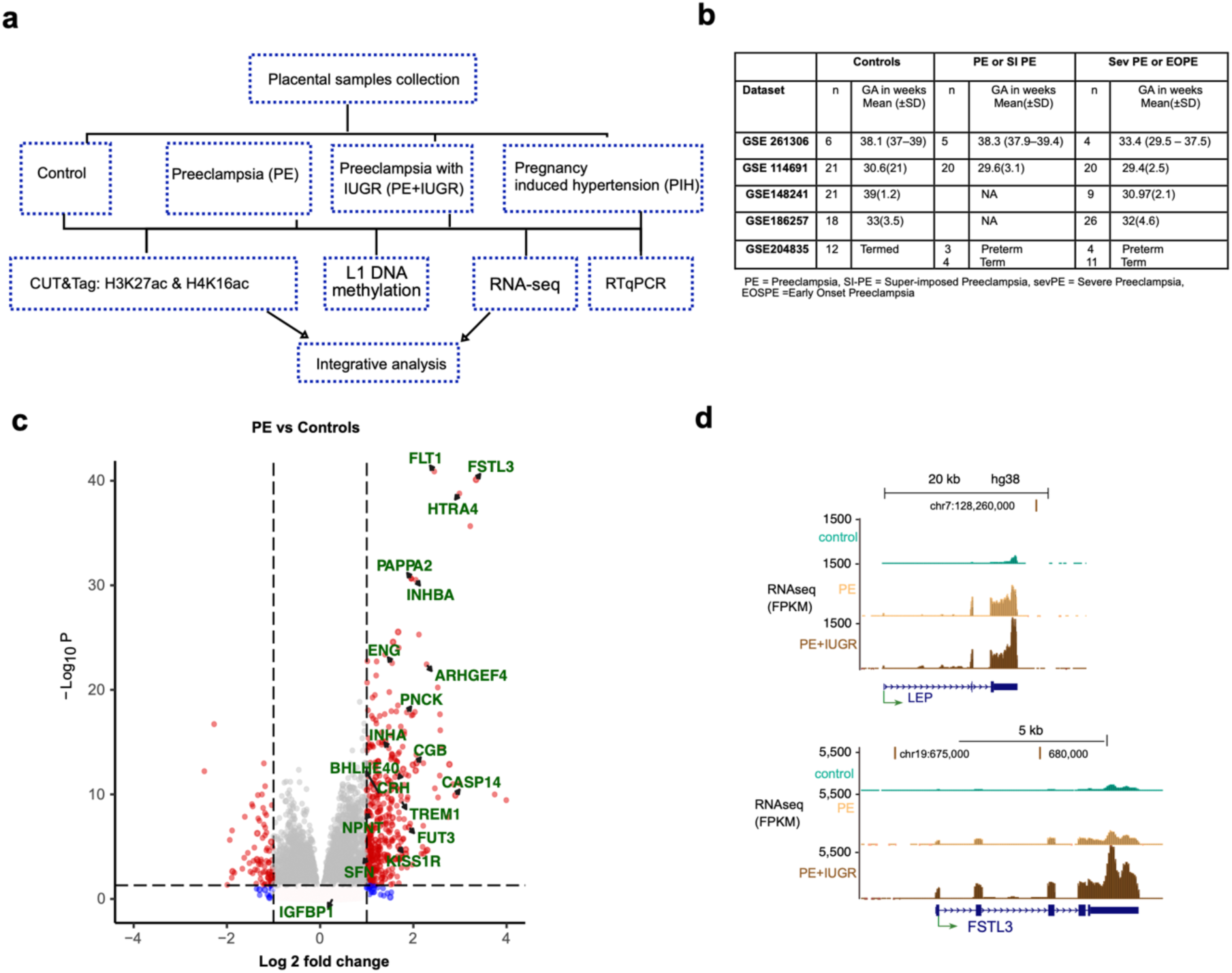
Genes and pathways dysregulated in PE. a) The study design for the cohort used in this study b) Placental biopsy cohorts analysed in this study with details of PE classifications, number of samples, and gestational age (GA) of the biopsies. c) Volcano plot showing significant (p adj or FDR <0.05) differentially expressed genes in PE vs healthy control placentas (list of DEGs in source data file 1). Genes known to be upregulated in the PE placenta are labelled. d) Genome-browser (hg38) snapshots showing average RNAseq reads across PE-associated LEP and FSTL3 genes in control (n=6), PE (n=5) and PE+IUGR (n=4) samples. Public placental RNAseq datasets used are listed in source data 1. DEGs for PE vs Control are mentioned in source data 2.

### TEs are deregulated in the preeclamptic placenta

ERV-derived genes, such as Syncytins and Suppressins, are essential for the development and functioning of the placenta ^50,51^. Moreover, many LTRs of ERVs are known to regulate placental-specific genes^52^. Hence, we aimed to identify differentially expressed TE transcripts in the PE placenta. We used the TE transcript tool to map sequencing reads at the subfamily level, which avoids bias against mapping to younger TE subfamilies ^53^, followed by differential expression analysis of TE subfamilies using DEseq2. Along with our placental biopsy data, we analysed differential expression from five independent cohorts, including two cohorts that contain gestational age-matched placental biopsies. Unsupervised clustering based on differentially expressed TE subfamilies (padj<0.05) from five independent cohorts led to a clear separation of PE from healthy control samples, irrespective of the disease severity and classification (PE, severe PE and PE with IUGR, superimposed PE) and timing of the onset of PE (early or late) (Fig. 2a). Separate analysis of gestational age-matched samples also showed similar pattern of TE expression changes in preeclampsia (Fig. 2b and Extended Data Figs. 2a and 2b) indicative of preeclampsia-specific differences in TE levels are not due to gestational age of the placenta. Further comparison of TE expression changes at individual loci levels showed upregulation of many L1 and ERV internal elements or LTRs are differentially expressed in the placentas with preeclampsia compared to controls (Fig. 2c, Extended Data Fig. 2c and 2d).

**Fig. 2.**
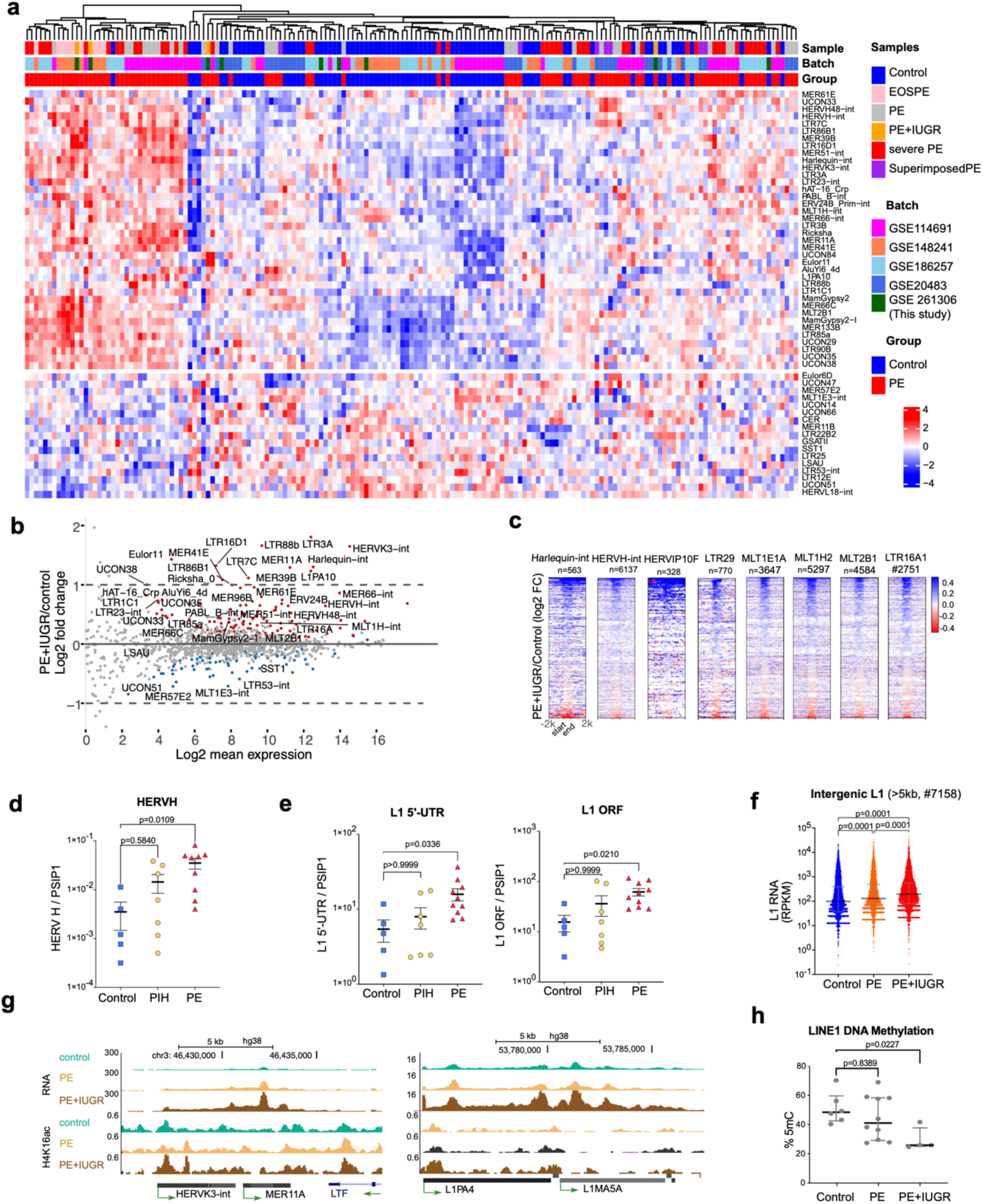
Multiple TE subfamilies are deregulated in preeclamptic placentas: a) Heatmap of the z-scores on Variance Stabilizing Transformation (VST) normalised counts of TE-subfamilies that are differentially expressed in preeclamptic placentas from five independent cohorts (Fig. 1b) with different PE conditions as annotated in each cohort (FDR p adj < 0.05, differentially expressed TEs listed in the additional data file 2). b) MA plot for differentially expressed TEs in PE+IUGR vs Controls for gestational age-matched controls from GSE114691. TEs consistently deregulated from panel 3a are labelled. c) Heatmap of RNAseq reads (log2 PE+IUGR/controls) for randomly chosen representative candidate TE-subfamilies across individual TE loci. d and e) RTqPCR quantification of HERVH and L1 5’-UTR and L1 ORF1 transcripts (normalised to PSIP1) in controls, pregnancy-induced hypertension (PIH) and PE (PE and PE+IUGR) placentas. f) Scattered dot plots comparing the expression of full-length intergenic L1s (as RPKM on the y-axis) across control, PE and PE+IUGR samples (p values indicate the significance of non-parametric Friedman test for multiple paired comparisons). g) Genome-browser snapshot of representative L1 and HERV LTR loci showing RNA (RPKM) and H4K16ac counts per million (CPM) levels. g) ELISA based L1 5-methyl Cytosine (5-mC, DNA-methylation) assay showing the percentage of L1 5’-UTR methylation between control, PE and PE+IUGR samples. p values for DNA methylation and RT-qPCR assays were calculated using ANOVA and the Kruskal-Wallis test for multiple comparisons. Differentially expressed TEs for PE vs Control are mentioned in source data 3 (for all cohorts integrative analysis) and 4 (for data generated in this study).

Many upregulated TE subfamilies in PE placentas included ERVs LTRs, DNA transposons and *Alu*s (p adj < 0.05 Figs. 2a and 2c and Extended Data Fig. 2e). Since L1s did not reach the significance threshold, we quantified their expression level separately, which revealed the upregulation of some L1s in PE samples with a low fold change (p adj < 0.05 Extended Data Fig. 2f). Upregulation of HERVs and L1s in PE is validated by RT-qPCR using primers that detect multiple loci of HERVH family and L1 5’ UTRs of evolutionarily young L1 elements (L1HS–L1PA5) and L1 ORF1 (Figs. 2d and 2e). Moreover, the changes in expression at these L1s were not an artefact of pervasive transcription as intergenic (for protein-coding genes) full-length L1s (>5kb) showed significantly higher expression in PE+IUGR compared to healthy controls (Fig. 2f and 2g). However, this RTqPCR validation showed no significant deregulation in pregnancy-induced hypertension without proteinuria (PIH, gestational hypertension), suggesting that the upregulation of TEs is specific to PE (Figs. 3c and 3d).

**Fig. 3.**
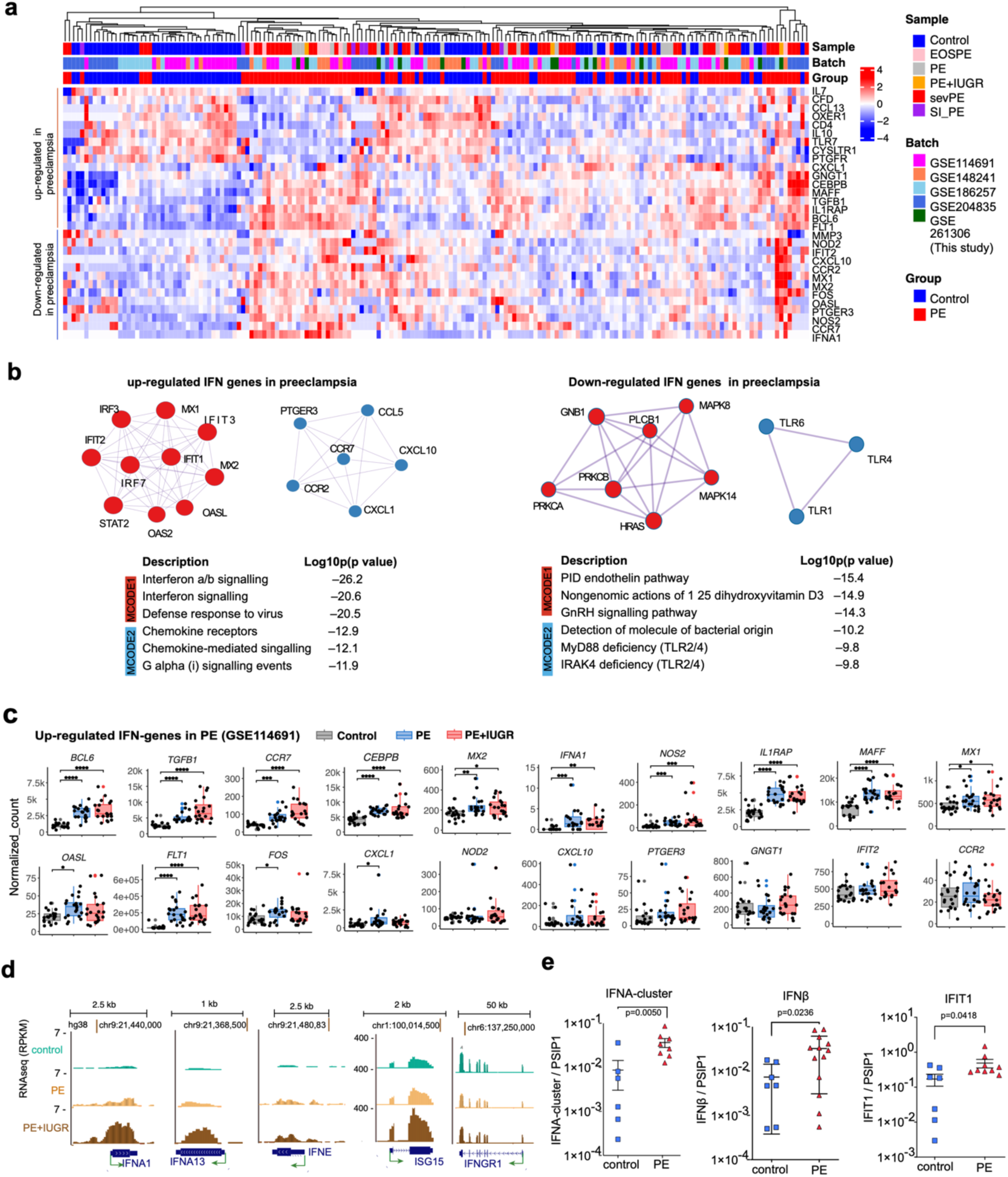
Type 1 IFN pathway is upregulated PE placenta. a) Like Fig 2a. but for differentially expressed IFN genes in Preeclampsia. b) Gene ontology network analysis (using metascape) of upregulated and downregulated IFN-related genes (p adj < 0.05). c) Box plots showing RNA levels for IFN genes upregulated in PE as in panel 3a for gestational age-matched GSE114691 cohort (p values represent Dunn’s test for multiple comparisons). d) Genome-browser track showing average RNAseq reads from control (n=6), PE (n=5), and PE+IUGR (n=4), each having two biological replicates at IFN-1 (IFNA1, IFNA13, IFNE, ISG15) and IFN-II (IFNGR1) gene loci. e) RT-qPCR in PE+IUGR and control samples for IFNß, IFIT1, and IFNA primers (Extended Data Table 2) that amplify multiple IFNA cluster genes. p values for RT-qPCR assays were calculated using ANOVA and the Kruskal-Wallis test for multiple comparisons.

### Elevated type-I interferon pathway in PE placenta

Due to sequence and structural similarities between TE-derived nucleic acids and viruses, cells sense TE transcripts and reverse transcribed cDNA as invading viruses, triggering the major antiviral IFN-I innate immune pathway ^9,54–56^. Hence, we aimed to test the hypothesis that higher TE transcript levels could lead to a higher IFN-I pathway. We probed thse known IFN-pathway genes (n= 250, from the Interferome database^57^) in the RNAseq data from multiple independent cohorts if they are differentially expressed in PE. Indeed, unsupervised clustering of differentially expressed IFN pathway genes showed clear separation of PE and control samples, particularly many IFN-I genes such as IFNA1, ISG15, MX1, MX2, and STAT2 were upregulated in PE samples (Fig. 3a). Pathway enrichment analysis of all upregulated IFN responsive genes showed enriched for IFN-I but not the IFN-II pathway, while downregulated IFN responsive genes did not show enrichment for IFN-I pathway (Fig. 3b). Furthermore, the quantifying expression level of IFN-I genes in a gestational age-matched cohort confirmed that upregulation of IFN-I pathway genes in PE is not due to lower gestational age in some PE placenta (Fig. 3c and Extended data Fig. 3a). Likewise, downregulated IFN pathway genes (Fig. 3a) showed lower expression levels in the gestational age-matched PE placenta than in healthy controls (Extended data Fig. 3b). RT-qPCR in our placental biopsies further confirmed the upregulation of IFN-I genes in PE placentas (Fig. 3d and 3e). Overall, we demonstrate that higher levels of L1 transcripts, ERV internal regions, and LTRs in the PE placenta correlate with elevated levels of the antiviral IFN-I pathway. Further experimentation using in-vitro TSC models is needed to establish the direct effect of deregulated TEs in a chronic inflammatory state associated with PE ^58^.

### H4K16 and H3K27 hyperacetylation in preeclamptic placentas

To elucidate the contribution of altered chromatin modifications at TEs and IFN-I deregulated in the preeclamptic placentas, we mapped genome-wide enrichment of acetylation of H4K16ac and H3K27ac by performing two replicates of CUT&Tag from different parts of placental biopsies from healthy controls (n=6) and PE+IUGR biopsies(n=4) (Fig. 1a and Extended Data Table 1). We evaluated the overall data quality and similarity among CUT&Tag data from independent placental samples and replicates (Extended data Fig. 4a). H3K27ac level was higher in genes upregulated in PE. However, the H4K16ac level was unaltered and showed poor correlation with the expression levels of upregulated genes (Fig. 4a, Extended data fig. 4b). This is consistent with recent findings from our lab and others demonstrating that MSL-mediated H4K16ac does not directly impact the expression of genes ^18,59,60^. Meta-analysis revealed the expected enrichment of H3K27ac at gene promoters or transcription start sites, but H4K16ac is enriched at gene bodies and enhancer features in PE and PE+IUGR compared to control (Fig. 4b and Extended data fig. 4a).

**Fig. 4:**
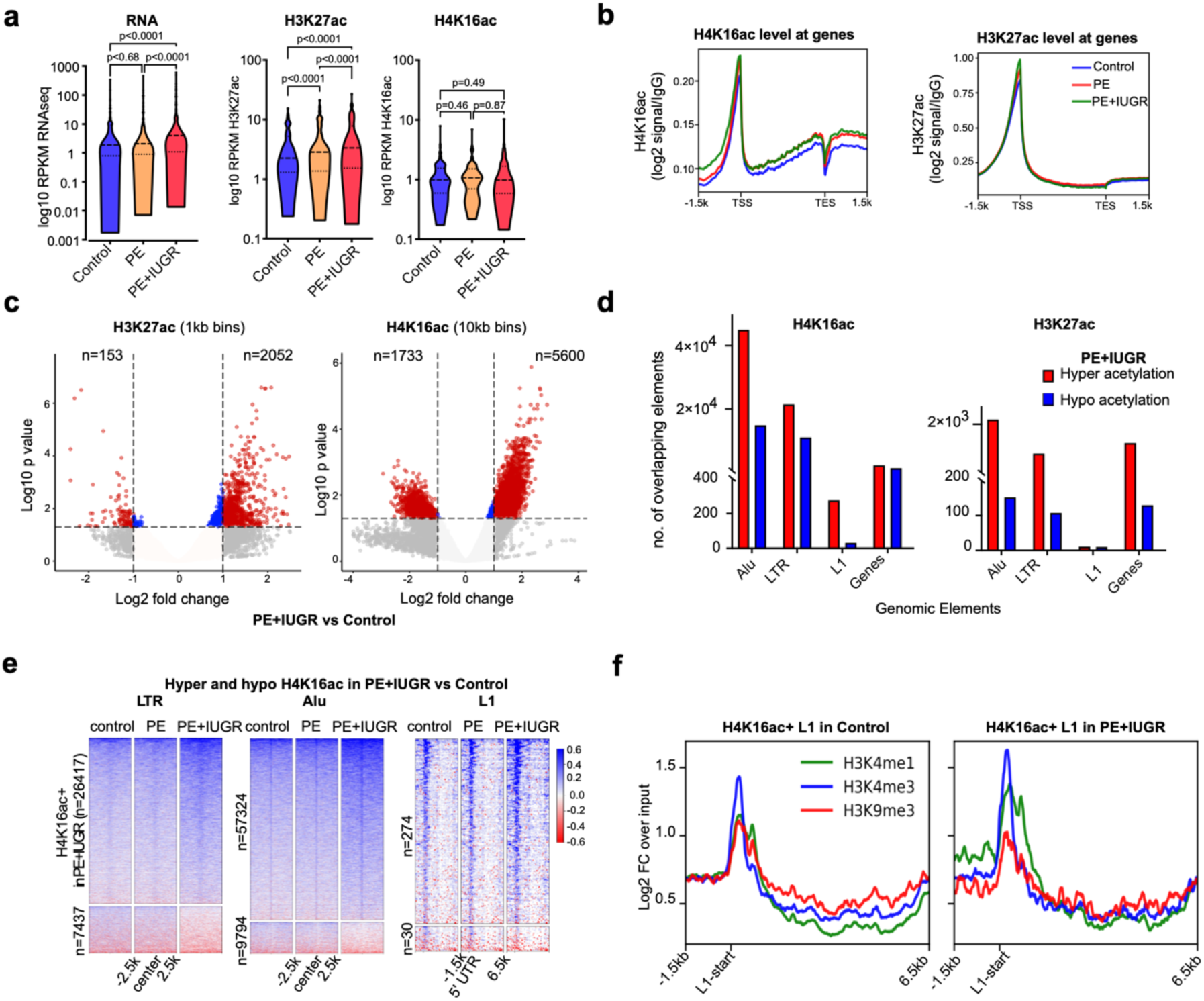
H4K16ac levels correlate with TE expression in PE. a) Violin plots for the RNAseq (RPKM), H3K27ac and H4K16ac CUT&Tag read counts per million (CPM) for the up-regulated genes (n = 88) in PE+IUGR compared to control. b) Meta profile for average H4K16ac and H3K27ac CUT&Tag reads in control, PE and PE+IUGR placentas across protein-coding genes (n=19291). c) Volcano plots showing significant (FDR <0.05, red) differential H4K16ac (10kb) and H3K27ac (1kb) genomic domains in PE+IUGR compared to healthy controls. d) Bar plots for the number of protein-coding genes (genes) along with full-length L1 (>5kb), Alu and LTR elements overlapping differentially acetylated H4K16ac (left) and H3K27ac (right) domains in PE (c). e) Heatmaps for IgG normalised H4K16ac signal across LTRs, Alus and full-length L1s overlapping hyper (top) and hypo (bottom) acetylated H4K16 domains in PE+IUGR compared to control. f) Metaplot for H3K4me1, H3K4me3 and H3K9me3 from chorionic villi at full-length LINE1s that are hyperacetylated in control (left) and in PE+IUGR (right). P values for violin plots were calculated using Friedman test for non-parametric paired ANOVA for multiple comparisons. List of CUT&Tag replicates are mentioned in source data 5. Differentially acetylated regions for H3K27ac and H4K16ac are listed in source data 6 and 7 respectively.

Further, to assess the changes in acetylation levels across the genomic features, we analysed the coverage of CUT&Tag reads across genomic bins of 1 kb with 0.5kb sliding windows for H3K27ac and 10 kb with 5kb sliding windows for H4K16ac due to its broad enrichment pattern ^59^. We detected more genomic bins enriched with H3K27ac and H4K16ac in the PE+IUGR placenta compared to control samples (Fig. 4c). H4K16 hyperacetylated domains showed overlap with enhancer marks such as H3K4me3, H3K4me1 while hypoacetylated H4K16ac domains overlapped with repressive marks such as H3K27me3 and H3K9me3 (Extended Data fig. 4c). Notably, H4K16ac domains specific to PE+IUGR are enriched at full-length L1s, *Alu* and LTR elements but not at genes. Whilst H3K27ac domains specific to PE+IUGR are enriched at genes, ERV LTRs and *Alu* elements but not at full-length L1s (Fig. 4d and e). L1s (>5kb) having higher H4K16ac levels showed relatively higher enhancer marks (H3K4me1 and H3K4e3) with low H3K9me3 levels at the 5’ UTR in PE+IUGR compared to healthy controls (Fig. 4f). Overlap of H4K16ac bins with L1s in PE & PE+IUGR suggests the contribution of H4K16ac to L1 expression levels in preeclampsia, which is consistent with our recent work, which demonstrated the role of H4K16ac in the transcription of L1s ^18^.

### L1s are hypo-methylated in PE placenta

A higher level of H4K16ac at L1 5’ UTRs at PE+IUGR (Fig. 4e) and significant upregulation of full-length L1s located outside the genes (Fig. 2f and 2g), confirming L1 upregulation is not due to readthrough transcription of introns of genes harbouring L1s. H4K16ac level increases globally with ageing and is associated with the loss of heterochromatin in senescent cells ^20,21^. Reduced heterochromatin and DNA methylation contribute to ageing and senescence-associated increases in TE transcription ^12^. DNA methylation plays a vital role in transcriptional repression of retroelements in somatic tissues. The reactivation of HERVs and L1s associated with reduced DNA methylation increases with age ^61,62^. Thus, we measured 5-methylCytosine (5-mC) across L1 repeats in placental biopsies. PE and PE+IUGR placentas showed hypomethylation at L1s compared to the control placenta (Fig. 2h). Since L1s constitute ∼18% of the human genome, hypomethylation at these elements in PE suggests possible global hypomethylation in PE, consistent with the previous findings demonstrating hypo-methylation signatures associated with PE ^33^. Our findings show that DNA hypomethylation, histone hyperacetylation and the upregulation of TE transcripts in the preeclamptic placenta further establish the epigenetic deregulation of TEs in PE. However, further work will be necessary to establish the impact of PE specific DNA hypo-methylation on TE expression and on placental development and function.

### Differential levels of TEs in cfRNA from preeclampsia pregnancies

The placenta is a fast-growing and transient organ, and necrosis and apoptosis of trophoblast cells are known to be associated with preeclampsia. The preeclamptic placenta is known to have high senescence and accelerated ageing phenotype plasma ^3,26,63^. This suggests that altered proliferation, senescence, and necrosis of placental cell types could contribute to the transcripts detected in the maternal plasma. Recent work shows placental cells undergoing necrosis and apoptosis are known to contribute significantly to circulating nucleic acids in the maternal blood, which can provide a non-invasive and early prognosis of PE ^30–32^. TEs are highly cell-type specific and are expressed at higher levels during early development, and in the placenta ^23,24^. Thus, we aimed to test whether we can detect deregulated TE transcripts in maternal cfRNA by performing differential expression analysis of TE subfamilies in cfRNAseq data from a prospective pregnancy cohort generated in Moufarrej et al. 2022 ^31^. Analysis of > 13-week gestational age (GA) cfRNAseq data showed a significant difference in the transcript levels of 133 TE subfamilies between healthy control and PE samples (*p* adj<0.05). Unsupervised clustering of samples based on these TE expression levels segregated PE from control samples (Fig. 5a and 5b). Many L1s subfamilies, including L1PA6, L1PA17, L1M, ERV LTRs subfamilies including MER11B, LTR25-int, LTR-12, HERVK9-int and HERV1-int and SINE including *AluSx4, Mir3* were detected at a significantly higher level in pregnancies that later developed PE (Fig. 5a and 5b).

**Fig. 5:**
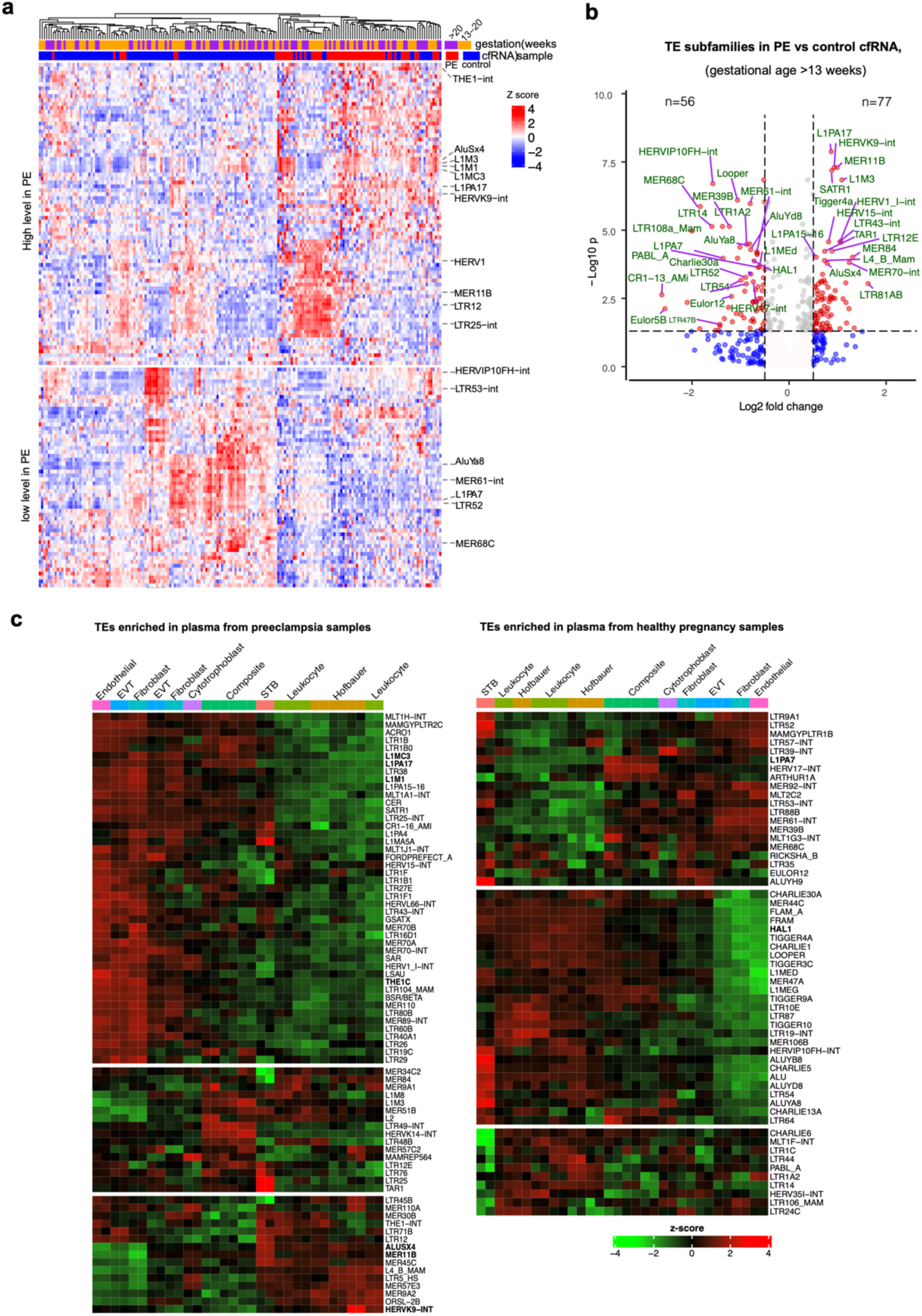
TEs transcript levels in maternal cfRNA. a) Heatmap for VST counts (z-scores) for 133 TE-subfamilies showed differential abundance between PE and control plasma RNA samples collected >13 weeks of gestation. b) Volcano plot showing significantly altered levels of TE subfamilies in the PE samples compared to control (FDR <0.05, list of deregulated TEs in the additional data file) c) Heatmap for RNAseq (VST counts) data from FACS sorted placental cell-types placental biopsies (GSE182381). TE subfamilies are significantly higher in PE plasma samples level than the control (left), and TE subfamilies are significantly lower in PE plasma compared to healthy controls (right). CfRNA datasets used for analysis in this study are listed in source data 8 and differentially abundant TEs in PE cfRNAs are listed in source data 9.

### Source of TE transcripts in maternal cfRNA

TEs are upregulated in early developmental stages, including embryonic stem cells and in the trophectoderm from which the placenta is derived ^23,24^. Notably, many TEs that show differential levels in cfRNA data, such as MER61, MER39B, MER11B, MER11A, MLT1, MLT2, MLT2B1 and L1 subfamilies are also differentially expressed in preeclamptic placental biopsies compared to healthy controls (Fig. 2a). We aimed to investigate the placental cell types contribution to cfRNA signature in the maternal cfRNA. For this purpose, we analysed the TE expression level in the RNAseq data from FACS-sorted placental cell types^64^. We found many TE subfamilies detected at higher levels in PE cfRNA compared to control are upregulated in cytotrophoblasts, EVTs, endothelial and fibroblasts compared to Hofbauer cells and Leukocytes in the placenta (Fig. 5c). This suggests a major contribution of trophoblast-derived cells to TE levels detected in the maternal cfRNA.

Although STBs also show upregulation of some TE subfamilies, TEs upregulated in the STBs differ from cytotrophoblasts and EVTs. Notably, TE subfamilies detected at lower levels in PE are downregulated in EVTs, endothelial, and fibroblasts compared to Hofbauer cells, leukocytes, and STBs (Fig 5d). HERVK9-int levels are consistent with the LTR5-Hs levels known to regulate the expression of the HERVK subfamily of internal regions. However, not all TEs detected at higher levels in PE cfRNA agree with this trend, as some TEs, such as MER11B, AluSx4 and LTR5-Hs, are detected at lower levels in EVTs but are upregulated in Hoffbauer, leukocytes and STBs. This could be due to the contribution of foetal and maternal organs to these cfRNA levels in maternal plasma ^65^.

### TE-cfRNA signature predicts PE

Two recent publications demonstrated that mRNA and noncoding RNA-based cfRNA signatures effectively predict who develops PE symptoms^66,67^. We used cfRNA datasets from Moufarrej et al. 2022 to train models that predict PE based on expression levels of TE subfamilies in cfRNA. For this purpose, we randomly split > GA13 samples into discovery (60 control and 29 PE cases) and validation (55 control and 31 PE cases) samples and compared the variance stabilised (VST) counts for predictions. We first performed feature reduction by applying a regularised random forest (RRF) algorithm on 133 differentially enriched TEs (Fig. 5a and 5b) for all samples >13 weeks GA. We selected 41 TE subfamilies with a variable importance of >0.5, unsupervised clustering based on VST normalised TE expression levels effectively segregated samples into control and PE (Extended Data Fig. 5a). Linear Discriminant Analysis (LDA) on these TEs to train the model resulted in a near-perfect AUC of 0.99 for the discovery and 0.77 for the validation samples (Extended Data Fig. 5b). Final signature after selecting 11 TEs based on linear-discriminant (LD) coefficients (Fig. 6a) and differential expression in discovery cohort (source data), improved the predictability with the AUC of 0.94 for discovery and 0.88 for validation samples (>13 weeks) with 81% sensitivity, 84% specificity, 74% positive predictor value (PPV) and 88% negative predictor value (NPV) (Figs. 6b and 6c, Extended Data Table 3 and 4).

**Fig. 6.**
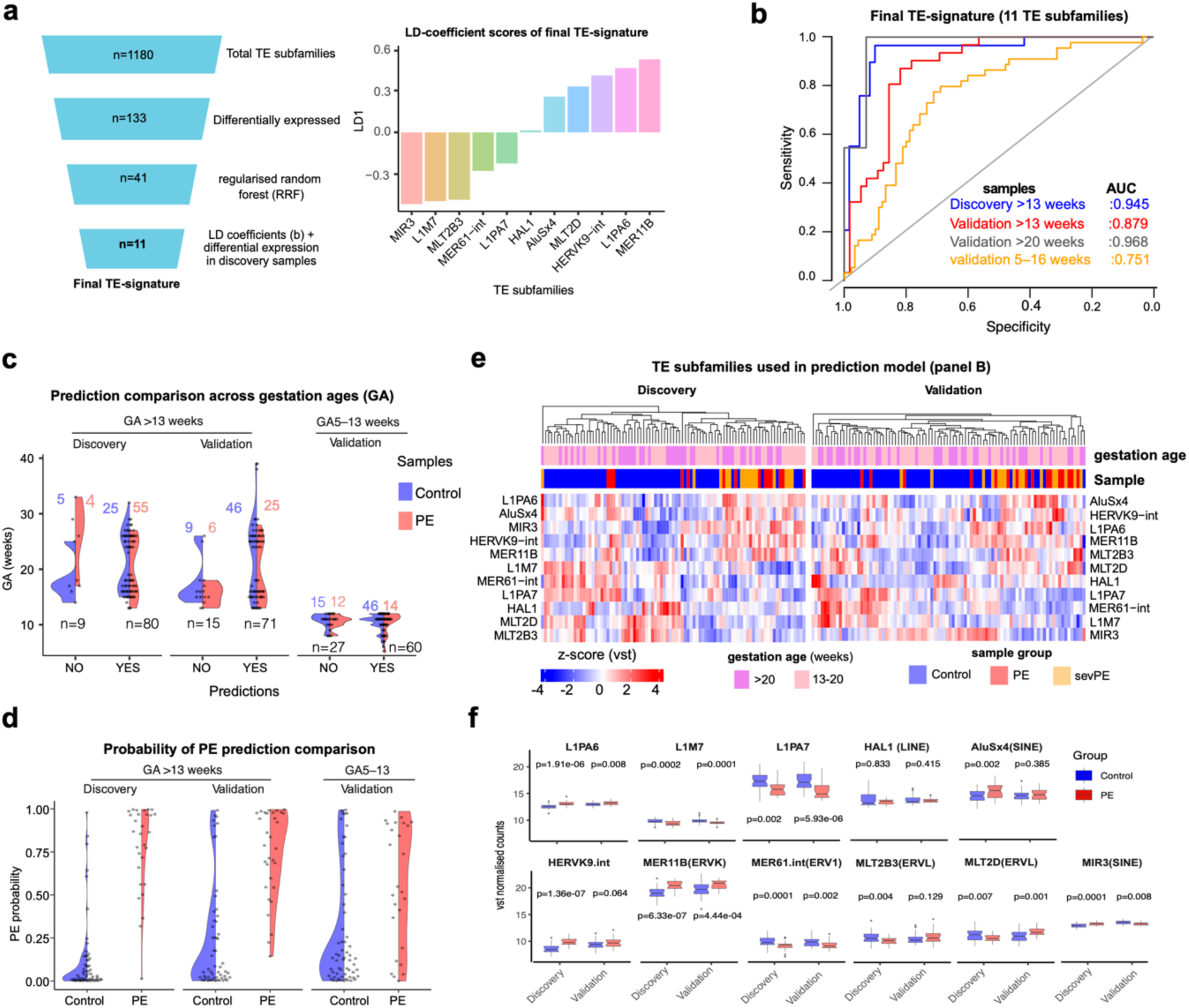
TE signature accurately predicts PE. *a)* Schematics (left) showing the training models used to identify the final TE signature that predicts PE (RRF variable importances for 133 TE-subfamilies listed in the additional data file). Linear Discriminant (LD) coefficients (y-axis) of the TEs (x-axis, n of TEs=11) used for LDA modelling (right). B) Area under the curve (AUC) with 95% confidence interval (right) for >13, >20 and 5–16 week gestation age (GA) showing sensitivity (y-axis) and specificity (x-axis) for PE prediction model using the final list of 11 TEs (Extended Data Table 3) for discovery (blue) and validation (red, grey and orange) cohort samples comprising different GA-groups. c) Split-violin plots showing the distribution of the predictions (x-axis) of indeed predicted (Yes) and falsely predicted (No) controls (blue) and PE (red) cases across the gestational ages (GA) (y-axis) for >13 weeks discovery, validation and GA 5–13 weeks samples using LDA model for TE signature from (b). d) Distribution of the probability of PE prediction values for the control and PE samples across the discovery, validation GA>13 weeks and GA 5–13 weeks samples for TE signature. e) Unsupervised clustering of discovery and validation samples based on expression level (VST counts) of TE signature for control and PE cfRNA samples with GA group (13-20 weeks and >20 weeks) and disease severity (PE, severe PE and control). f) Box plots comparing normalised RNA counts for final TE subfamilies used in (b) for the discovery and validation cohort (GA>13 weeks). p values were calculated using Dunn’s test for multiple comparisons. RRF variable importances for 133 TE subfamilies are listed in source data 10. differentially abundant TEs in PE cfRNAs for discovery and validation cohorts are listed in source data 11 and 12 respectively.

Moreover, re-running the predictions for validation cohort with gestational age group similar as in Moufarrej et al. 2022 (GA 5-16 weeks) showed effective at distinguishing between control and PE with AUC 0.75 (Fig. 6c and Extended Data Fig 5b, Extended Data Table 4) indicating synergistic contribution of these TE levels to predict PE. Compared to the recent gene expression signature for early GA (5–16 weeks) that showed 28% PPV and 71% NPV ^31^, our TE signature in validation samples of the same GA showed 57% PPV and 81% NPV (Extended Data Table 4). This TE-based prediction scores are superior to previous mRNA- and noncoding RNA-based predictions that showed 75% sensitivity and 32.3% PPV^31^. Moreover, this signature showed increased AUC values when compared for prediction across different gestational age groups (>13 weeks) (Fig 6b and 6c and Extended Data Fig 5c). Interestingly, further sub-set of 6 and 3 TEs showed an AUC of 0.83 in the validation cohort (GA >13 weeks), suggesting that PE prediction can be achieved using small TE subfamilies (Extended Data Fig. 5d and Extended Data Table 4).

The identified TE-cfRNA signature showed consistent predictability of PE cases with a higher probability distribution (>0.5) for predicted PE cases compared to controls (<0.5) (Fig. 6d). Expression level of this final TE signature was sufficient to segregate >13 weeks GA control and PE samples irrespective of the GA and PE severity (Fig. 6e). Five TE subfamilies showed a same trend and significant difference in the TE levels between control and PE samples between discovery and validation cohort. Two TEs showed the same trend in the validation cohort but did not reach a significance level (p<0.05, Dunn’s test) (Fig. 6f). Our analysis demonstrates that TE transcript levels can predict PE cases early in pregnancy independent of the GA and PE severity. We show that TE transcript levels can be used as a new non-invasive approach for predicting PE cases early in pregnancy independent of PE severity. Adding maternal and medical data together with this non-invasive approach to predicting PE early in the pregnancy will allow prophylaxis at an early stage, improving pregnancy outcomes.

## Discussion

The majority of pregnancy complications, including PE, IUGR, premature birth and stillbirth, are associated with placentation defects and inflammation. Although inflammation during pregnancy is a major cause of pregnancy complications leading to long-term effects on both mother and child, it is unclear what causes inflammation during pregnancy. Here, we demonstrate that TE deregulation associated with H4K16ac hyperacetylation leads to the activation of the antiviral IFN-I pathway, suggesting the role of deregulated TEs in inflammation in preeclamptic placentas. Using these insights, we have discovered cfRNA TE signatures that can be utilised for early diagnosis of pregnant women at risk of developing PE early. This improves targeted surveillance and obstetric care.

PE-specific hyperacetylation at TEs leading to their upregulation could contribute to the preeclamptic placenta, as accelerated ageing-like state and senescence are implicated in the pathophysiology of PE ^3,26,63^. We found a higher level of acetylation in PE placenta with IUGR but not in late-onset PE, suggesting an altered level of acetylation could be linked to different pathophysiology as it is known that early- and late-onset PE have different pathophysiology ^68^. H4K16ac is related to ageing and senescence ^19,21^, suggesting the contribution of TE-specific hyperacetylation to the PE phenotype. What causes altered H4K16ac levels in PE is not clear. We hypothesise that the altered glycolysis and hypoxia pathways (Extended Data Fig. 1d) could contribute to acetylation due to altered acetyl-CoA levels in the PE placenta. The contribution of diet-induced changes in acetyl-CoA in altered levels of histone acetylation at TEs also cannot be ruled out, as the higher levels of nucleocytoplasmic acetyl-CoA can serve as a substrate for histone acetylation in growth or fed conditions compared to starved conditions (reviewed in) ^69^. This agrees with the hypothesis that chromatin modifications enriched at TEs, constituting nearly 50% of the genome, can be a source or sink for metabolic by-products such as acetyl-CoA (discussed in) ^70^.

TEs are known to be upregulated during early development, embryonic stem cells and trophectoderm-derived trophoblast stem cells in the placenta ^22–24^. Trophectoderm gained repressive H3K9me3 domains are preferentially deposited at hominoid specific TEs such as LTR12, MER11B, HERVH, and HERVK9-int that are differentially enriched in PE cfRNA and placenta ^71^. PE is a human-specific disorder. Epigenetic remodelling of hominoid-specific TEs, which have co-opted placental-specific functions, is essential for early development, and their deregulation can lead to defective placentation or embryo implantation ^23^. Many LTRs such as MER84, MER61E, MLT1 MER11, LTR12, MLT1J2 and MER39B that are differentially expressed in cfRNA are suggested to function as cis-regulatory elements for placental or trophoblast-specific genes ^24,42,43^, supporting the contribution of placental cell types to maternal cfRNA signature. Increasing PE prediction accuracy for >20 weeks GA with 83% PPV, 0.96 AUC and 0.91 sensitivity compared to 5-16 weeks GA with 65% PPV, 0.75 AUC (Fig 6b, Extended Data Table 4) further supports the increased placental contribution to TE-cfRNA signature with the gestation ^72^.

Preeclamptic placental biopsies are significantly enriched with EVTs and depleted of STBs ^73^. Early onset PE is associated with a smaller birth and placental weight ^74^. This suggests an altered cellular composition of placental cell types in early-onset PE due to senescence or necrosis can contribute to TEs-cfRNA levels. A higher level of IFITM (an IFN-1-regulated gene) is shown to inhibit cell fusion in STBs and has been suggested to contribute to pregnancy complications ^75^. Our findings demonstrate that elevated IFN-1 pathways and reduced STBs in PE placentas suggest a possible role of the IFN-I pathway in altered trophoblast differentiation. Recent work shows an interesting link between pregnancy-specific upregulation of HERVs and elevated IFN-I response, and this TE-IFN-I axis is co-opted to activate haematopoietic stem cells and erythropoiesis ^76^. Further work is needed to establish a causal effect of the TE-IFN-I axis in placental development and function and their role in preeclampsia.

In summary, TE expression signatures identified here will provide accurate and non-invasive methods to predict PE early in pregnancy. The current PE screening models are based on maternal characteristics and medical history or on specialised ultrasound, blood analyte and hypertension measurements, which are insufficient for effective risk prediction. Our TE signature with 83% PPV is superior to the sFlt-1:PlGF ratio test currently recommended for 20 to 35 weeks GA with suspected PE, which provides a PPV of 36.7% ^77^. TEs cfRNA-based prediction also performs better than the non-TE RNA-based prediction approach ^31,32^. Further work to select a better combination of TE and non-TE transcript signatures and clinical and laboratory data collected during routine antenatal visits can improve PE prediction. The TE-cfRNA-based approach of predictability of PE across the gestational ages can be used for stratification of pregnancies for close monitoring and therapeutic interventions that may improve maternal and neonatal outcomes.

## Limitations of the study

One limitation of analysing the bulk epigenome and transcriptome of placental biopsies is the confounding effect of the placenta’s complex cell types. Thus, in-vitro experimentation is needed to decipher the direct impact of acetylation-mediated TE deregulation and its effect on triggering the IFN-I pathway using the trophoblast stem cell model.

## Acknowledgements

We thank patient families for participating in the PE epigenetics study and donating placental biopsies. We also thank members of the Madapura lab and QMUL Epigenetics Centre for discussions and reagents and Sarah Teichmann and Ioannis Sarropoulos (Sanger Institute) for their discussions and comments on the manuscript. This research used Apocrita HPC, supported by QMUL Research-IT.

## Funding

Medical Research Council UKRI/MRC grant (MR/T000783/1) (MMP, MP, FB), Barts charity small grant (MGU0475) (MMP). Newton Mosharafa scholarship from the British Council, Egypt, and the Central Department of Missions, Egypt (AS, ASA).

For Open Access, the author has applied a CC BY public copyright license to any Author Accepted Manuscript version arising from this submission.

## Author contributions

MMP, MP, AS and ASA acquired the funding, conceived and designed the study, and supervised the work. AS wrote the PE epigenetic study protocol and obtained ethical approval. ASA recruited the subjects and collected placental tissue samples and clinical data. MP and ASA performed the CUT&Tag, RNAseq experiments, FB and AD performed RTqPCR with contributions from MMP, and MP analysed CUT&Tag, ChIPseq, bulk RNAseq and cfRNAseq data. MMP and MP wrote the manuscript. All the authors have read and approved the final version of the manuscript.

## Materials and Methods

### Ethics statement

Patients were enrolled at the Royal London Hospital, Barts Health Trust from May 2021 to March 2022 as a part of a PE Epigenetics study with written informed consent before participating and ethical committee approval (REC 21/SS/0010) from the UK Health Research Authority. Demographic and clinical details were obtained from the Clinical Record Service (CRS) Cerner Millennium and BadgerNet Clevermed database. Immediately after birth, we collected the samples from the fresh placenta. Four samples (1cmΧ 1cm) of villous tissue were accessed by trimming away the basal plate with a pair of scissors and a scalpel, then cutting out a piece of the exposed villous tissue and discarding the basal plate tissue. Dissected villous tissue was immediately transferred to a dish containing PBS, and the tissue was cut into small pieces, then snap-frozen immediately and stored at -80°C. We performed two replicates of CUT&Tag for H3K27ac and H4K16ac and polyA-RNAseq from different parts of the placenta biopsies. Sequencing data that failed were excluded for further analysis.

### Recruitment criteria

Babies born to women with PE were defined as recommendations from the International Society for the Study of Hypertension in Pregnancy (ISSHP) ^78^, PIH, and normotensive control who delivered at the Royal London Hospital and were willing to provide informed consent. Exclusion criteria: Infants who were critically ill and babies with significant congenital and genetic abnormalities. Three groups were recruited, normotensive control, PE with GA> 37 weeks without IUGR, and PE <37 weeks and IUGR. IUGR is defined as weight for GA < 5th centile. The last group contain four preterm pregnancies < 37 weeks and one pregnancy > 37 but with IUGR (Extended Table 1).

### RNA isolation and RT-qPCR

Total RNA from Placental biopsies RNA using TRI reagent solution (ThermoFisher Scientific, AM9738), genomic DNA was eliminated by treating RNA samples with Turbo RNAse free DNAse1 (ThermoFisher Scientific AM1907). For reverse transcriptase-polymerase chain reaction (RT-qPCR), cDNAs were prepared with LunaScript^®^ RT SuperMix Kit (NEB, E3010). qPCR was performed using qPCRBIO SyGreen Mix Lo-ROX (PCRBio) in LightCycler 480 instrument (Roche). Primer pairs for human L1 5’ UTR and L1 ORF1 to human L1s were designed to amplify elements of the human-specific L1HS preferentially and evolutionarily recent primate-specific L1PA (L1PA2–L1PA6) subfamilies were taken from ^61^. Primers to the human IFNA family against a consensus sequence of all human IFNA gene sequences

(*IFNA1*, *IFNA2*, *IFNA4*, *IFNA5*, *IFNA6*, *IFNA7*, *IFNA8*, *IFNA10*, *IFNA13*, *IFNA14*, *IFNA16*, *IFNA17* and *IFNA21*) and IFNB1 are taken from ^61^. The list of all primers used for RTqPCR is in Extended Data Table 2. Data were normalised to β-actin or PSIP1.

### RNA sequencing

RNA was isolated from placental biopsies using TRI reagent solution (ThermoFisher Scientific, AM9738), and genomic DNA was eliminated by treating RNA samples with Turbo RNAse free DNAse1 (ThermoFisher Scientific AM1907). RNA sequencing library preparation using NEBNext^®^ Ultra^™^ II Directional RNA Library Prep Kit for Illumina^®^ (NEB #E7765), followed by libraries, were sequenced as 150 bp paired-end reads using Novaseq 6000.

### LINE1 DNA methylation assay

Genomic DNA from placental tissues was isolated using a Quick-DNA mini prep plus kit according to the manufacturer’s instructions (Zymo Research D4068). LINE-1 methylation levels were quantified using an ELISA-based Global DNA Methylation Assay LINE-1 kit; the assay was performed as described by the manufacturer (Active Motif cat. no. 55017). Briefly, genomic DNA from each sample was digested overnight with MseI enzyme (10 U/μL) at 37°C. 100 ng of digested gDNA was hybridised with a LINE-1 probe in a thermal cycler (98 °C for 10 min, 68 °C for 1 hr, followed by a quick ramp to 25 °C). LINE-1 probe is a 5’ biotinylated oligo designed to hybridise to a 290 bp region of the LINE-1 repeat element, containing 88 cytosine residues, of which 12 are in a CpG context. Reactions were performed in triplicate along with the methylated and non-methylated DNA standard samples, prepared in parallel with placental genomic DNA samples. Digested DNA was transferred to a streptavidin-coated plate and incubated for 1 h at room temperature with mild agitation. Then, a 1:100 dilution of 5-methylcytosine monoclonal antibody was incubated for 1 hr at room temperature, followed by 1 hr of HRP-conjugated secondary antibody. The developing solution was added and incubated for 3 min; the stop solution was added when the standard samples showed colour change. Finally, the plate was read at 450 nm and 655 nm.

### CUT&Tag

CUT&Tag from placental biopsies was performed according to the Steve Henikoff lab protocol^79^, with modifications to tissue processing as described below. Different parts of placental biopsies were processed to perform replicates of H3K27ac and H4K16ac CUT&Tag. To adapt CUT&Tag tissue sections, flash-frozen placental tissues (approximately 3–4 mm size) were manually homogenised with tight homogenisers in wash buffer (20 mM HEPES pH 7.5, 150 mM NaCl, 0.1% BSA, 0.5 mM Spermidine and cOmplete EDTA-free protease inhibitor tablet) into a homogenous suspension of intact cells. Cells were transferred to 1.5-ml low DNA binding tubes (Eppendorf), and solutions were exchanged on a magnetic stand (DynaMag-2, Thermo Fisher Scientific). Cells were pelleted by centrifugation for 3 min 600×*g* at room temperature and resuspended in 500 μl of ice-cold NE1 buffer (20 mM HEPES-KOH pH 7.9, 10 mM KCl, 0.5 mM spermidine, 1% Triton X-100, and 20% glycerol and cOmplete EDTA-free protease inhibitor tablet) and let it sit for 10 min on ice. Nuclei were pelleted by centrifugation for 4 min 1300×*g* at 4 °C and resuspended in 500 μl of wash buffer, and the wash buffer by placing the tubes on a magnet stand to clear and withdraw the liquid, then resuspended in 1.0 ml wash buffer and held on ice until beads are ready. In total, 10 μl of BioMag Plus Concanavalin-A-conjugated magnetic beads (ConA beads, Polysciences, Inc) in binding buffer (20 mM HEPES-KOH pH 7.9, 10 mM KCl, 1 mM CaCl_2_, and 1 mM MnCl_2_) was added to each tube containing cells and rotated on an end-to-end rotator for 10 min. After a quick spin to remove liquid from the cap, tubes were placed on a magnet stand to clear and withdraw the liquid, and 800 μl of antibody buffer containing 1 μl of primary antibodies (normal rabbit IgG, Santa Cruz Cat no sc-2027, H3K27ac (Abcam, ab4729), H4K16ac (Abcam, ab109463) was added and incubated at 4 °C overnight in a nutator. Secondary antibodies (guinea pig α-rabbit antibody, Antibodies online cat. no. ABIN101961) were added 1:100 in Dig-wash buffer (5% digitonin in wash buffer) and squirt in 100 μl per sample while gently vortexing to allow the solution to dislodge the beads from the sides and incubated for 60 min on a nutator. Unbound antibodies were washed in 1 ml of Dig-wash buffer for a total of three times. In total, 100 μl of (1:250 diluted) protein-A-Tn5 loaded with adapters in Dig-300 buffer (20 mM HEPES pH 7.5, 300 mM NaCl, 0.5 mM spermidine with Roche cOmplete EDTA-free protease inhibitor) was placed on a nutator for 1 hr and washed three times in 1 ml of Dig-300 buffer to remove unbound pA-Tn5. Then, 300 μl tagmentation buffer (Dig-300 buffer + 5 mM MgCl_2_) was added while gently vortexing and incubated at 37 °C for 1 hr on an incubator. Tagmentation was stopped by adding 10 μl 0.5 M EDTA, 3 μl 10% SDS, and 2.5 μl 20 mg/ml Proteinase K to each sample. All were mixed by full-speed vortexing for ∼ 2 s and incubated for 1 h at 55 °C to digest. DNA was purified by phenol: chloroform extraction using phase lock tubes followed by ethanol precipitation. Libraries were prepared using NEBNext HiFi 2x PCR Master mix (Cat number M0541S) with a 72 °C gap filling step followed by 13 cycles of PCR with 10-s combined annealing and extension for enrichment of short DNA fragments. Pooled libraries were run on 1.5% ultrapure agarose gel, 200-700bp smear was excised, and DNA was extracted using Monarch gel extraction kit (NEB cat. No. T1020). Libraries were sequenced in Novaseq 6000 with 150bp paired-end reads at the Novogene sequencing service.

## Analysis of CUT&Tag data

### Mapping

150bp paired-end reads for the CUT&Tag-seq were trimmed for adapters using the Trimmomatic tool and aligned locally to the hg38 genome through Bowtie2 (version 2.4.5) with these parameters for pair-end mapping: *--very-sensitive-local --no-unal --no-mixed --no-discordant --phred33 -I 10 -X 700* ^80^. The best alignment was retained using default bowtie2 options for multiple aligned reads. The bam files were sorted, indexed, and used to generate bigwigs for individual replicates of H3K27ac and H4K16ac. Merged bam files were obtained across control, PE and PE+IUGR using *samtools merge* ^81^. These bam files were then sorted, followed by indexing and generating bed and bigwigs for individual modifications.

### Histone acetylation domain analysis

Counts were obtained for the H4K16ac and H3K27ac for either 10kb with a sliding window of 5kb (for H4K16ac) or 1kb with 500bp sliding window (for H3K27ac) genomic bins on hg38 genome across Control, PE and PE+IUGR groups using *bedtools multicov* tool. Domains containing less than 50 reads sum across all samples were filtered out from analysis for H4K16ac, and non-zero counts were used for H3K27ac analysis. DESeq was performed on these counts and plotted using the r-package *Enhanced Volcano* for the differentially acetylated regions (DARs). The threshold for the DARs was set at padj or FDR <=0.05 for Benjamini-Hochberg correction. The number of genomic elements (protein-coding genes, LTRs, Alu and full-length L1 >= 5kb) overlapping with the differentially acetylated regions were obtained for H4K16ac and H3K27ac respectively, using the *bedtools intersect* tool. Overlapping histone modification profiles were generated by submitting the DARs to the Cistrome browser ^82^ for H4K16ac and H3K27ac, respectively.

### Bigwig generation and plotting

Sorted bam files were subjected to bigwig generation via deepTools (version 3.5.1) ^83^ bamCoverage tool with *--binSize 20 –normalizeUsing CPM --scaleFactor=1.0 -- smoothLength 60 --extendReads 150 --centerReads* options. The signal was normalised to IgG through bigwigCompare. The bigwig files were used for plotting signals or visualisation in the genome browser. The genome-browser views were obtained by viewing the signal tracks in the UCSC genome browser or IGV.

Signal plotting at various genomic landmarks and bed coordinates was done using *deepTools*. Matrices were generated using deepTools *computeMatrix reference-point* or *scale-regions* option. These matrices were used for plotting heatmaps or average summary plots by *plotHeatmap* or *plotProfile* function in deepTools. Further comparisons on the hyper-or hypo-acetylated (H4K16ac) TEs for their potential as enhancers were confirmed by comparing the H3K4me1 (ENCFF710ASO), H3K4me3 (ENCFF169MHR) and H3K9me3 (ENCFF541CWH) available from the ENCODE datasets for Chorion villi.

### RNA-seq data analysis

The reads obtained from public placental RNAseq datasets (Fig. 1b) and our cohorts (two biological replicates per sample) were mapped to the human genome using STAR^84^. The bam files were merged for the replicates, followed by indexing and bigwig generation using the tools described for CUT&Tag samples earlier, with *normalisation using* RPKM. The counts for the genes were obtained using the *featurecounts* tool from the SubRead package, and TE counts at the subfamily were obtained using the TEtranscript tool with default options. The count matrices (genes or TEs) were subjected to DESeq2 for the differential analysis with default options. The PCA plot was generated using *plotPCA* function on rlog transformed data for RNAseq data generated in this study. While doing an integrative analysis of the public and our cohorts, the batch effect removal was performed for the differences in cohorts (each cohort serving as a batch) using *limma::removeBatchEffect* on the variance stabilised counts (VST) transformed count matrix. Differentially expressed genes (DEGs) or differentially expressed TE subfamilies were counted as those having Benjamini-Hochberg corrected FDR (padj) <=0.05. The DEGs were functionally annotated using *EnrichR* and *Metascape* for combined analysis. For our cohort, the functional enrichment was performed using *clusterProfiler (PMID: 22455463).* The differential expression of DEGs and TE-subfamilies was visualised as volcano plots or heat maps using EnhancedVolcano or pheatmap packages, respectively. For the heatmap, z-scores obtained on VST counts were used to compare the Control and PE cases.

For differential enrichment analysis at individual TE element level, the fragment counts for each dataset were obtained using the feature *counts* tool from the SubRead package for different TE classes (L1 and LTR), with the gtf fetched from the UCSC table browser. These feature counts were used for the differential enrichment analyses using the *DESeq2* package in R. The differential expression of genes or TEs was visualised as a volcano plot. For analysis of interferon-regulated genes, IFN-regulated gene lists were downloaded from the Interferome database ^85^. For comparison as a heatmap or functional annotation, only genes which are significantly dysregulated (p adj <0.05) for PE vs Control (for all cohort combined analysis) comparison were used. Functional annotation was done using *Metascape*.

The RNA signal across the intergenic (for protein-coding genes) full-length L1s (>5kb) was calculated as RPKM from the read counts obtained across the Control and PE+IUGR samples. This RPKM signal was then plotted as a scattered dot plot using GraphPad Prism 10. The signal was plotted as the *log10* value of the RPKM on the Y-axis. Paired violin-plot comparisons were compared using the ANOVA Friedman test for multiple paired data.

The RNAseq data for the cell-free (cfRNA) or plasma RNA was obtained from NCBI for the GEO series GSE192902 for the samples (Source data table). The reads were mapped onto the hg38 genome following a similar pipeline as for the placental RNAseq data described earlier. The counts for the TE-subfamilies were obtained using the TEtranscript tool with default options using gtf for genes and repeats (rmsk) from *ENSEMBL*. The TE count matrix was used to perform DESeq analysis. The comparisons were made within gestational age groups: GA <13 weeks, GA 13-20 weeks, and GA >20 weeks for either PE vs control or sevPE vs control. We initially compared the differential TE presence for PE (sevPE, also annotated as PE) vs Control across samples having gestational age >13 weeks. We obtained 133 TE-subfamilies which have significant (p adj <= 0.05, log2FC > or < ± 0.5) differential presence in PE cases (77 upregulated and 56 downregulated) as seen in the volcano plot (Fig. 5b). We plotted these individual TE subfamilies as a heatmap for all samples used in comparison using pheatmap tool, with k-means clustering on the z-scores obtained for the VST normalised counts. DESeq2 analysis was also performed for discovery and validation cohorts and for GA <13 and GA 5-16 weeks for further downstream usage in model signature selection and predictions.

### Cell-type specific comparison for differentially abundant TE subfamilies in PE-cfRNA

To deconvolute the placental cell types contribution to the cfRNA, we compared the levels of differentially abundant TE-subfamilies in PE-cfRNA across multiple placental cell types obtained by FACS-sorted placental biopsies. The cell-type specific bulk-RNAseq data was downloaded for GSE182381, followed by mapping and obtaining TE-counts as mentioned for Placental RNA-seq data. These TE counts were used to obtain the variance stabilise count using iDEP portal for RNAseq analysis. Comparison of VST counts across 133 differentially abundant TEs in PE-cfRNA was plotted as a heatmap of z-scores across placental cell types Leukocytes, Fibroblasts, Endothelial cells, Hofbauer cells, Syncytiotrophoblasts (STBs), Extra-villous trophoblasts (EVTs) and Cytotrophoblasts as well as composite placentas.

### Prediction model training and validation

We began with feature reduction on the VST normalised counts of significant differentially present TE-subfamilies in PE cfRNA (n=133) by applying the RRF (Regularised Random Forest) algorithm. Variable importance obtained by RRF for all 133 differentially abundant TEs are listed in the additional data file. We selected the 41 TE-subfamilies (variable importance >0.5) to model using linear discriminant analysis (LDA). We further used a combination of RRF variable importance and the differential presence of TEs in the discovery cohort to narrow the number of TE-subfamilies to 11 and further model using LDA. The models were trained on the Discovery cohort and tested on the validation sets (validation 1, GA <13 weeks, GA 5-16 weeks, 13-20 weeks, and GA >20 weeks) using LDA modelling. The AUC values were obtained using the pROC package, with confidence intervals (DeLong) of 95% calculated using *ci.AUC.* The following matrices were generated using the predictions obtained for the TE-subfamilies as listed in (Source Table)

1) sensitivity = TP/(TP + FN); (2) specificity = TN/(TN + FP); (3) PPV = TP/(TP + FP); (4) NPV = TN/(TN + FN), 5) accuracy = correct predictions (TP + TN)/ all predictions (TP+FP+TN+FN) TP =True Positive, FP = False Positive, TN = True Negative and FN = False Negative. Comparison of the probabilities to predict PE were used to plot between Control and PE for discovery and validation cohorts (validation 1 and GA 5-13 weeks) using the posterior values for PE for the LDA model trained or tested on them, respectively. The split violin plots were plotted using ggplot2.

### Statistical tests

For box-plots, and comparison of differentially expressed TE-subfamilies, the *Dunn test* function in the R tool *rstatix* with Bonferroni correction was used for multiple-group comparisons between the groups. All DESeq2 output was filtered for either FDR/ Benjamini-Hochberg adjusted p-value (padj < 0.05) or Wald test p-value (p < 0.05), as mentioned in the figure legends. P-values for RT-qPCR assays and gene-distance box plots were calculated using the ANOVA Kruskal-Wallis test for multiple comparisons using GraphPad Prism10. Paired violin-plot comparisons were compared using the ANOVA Friedman test for multiple paired data.

## Data availability

The data discussed in this publication have been deposited in NCBI’s Gene Expression Omnibus (GEO) and are accessible through the GEO Series accession number XXXX

CUT&Tag raw data and processed data files (bigwigs) can be accessed at NCBI with an accession ID, and RNAseq raw data files can be accessed with an accession ID. All the datasets generated and public datasets used in this study are detailed in (Source Table). CUT&Tag and RNAseq were performed using two different parts of the placental biopsies; replicates that failed QC were not used for the analysis.

## Code availability

All the analyses in this manuscript have been carried out using publicly available tools. No custom code was generated for this purpose. The methodology contains the details of the steps involved in the analysis.

