## Supplementary figures and tables for "Early Prediction of Preeclampsia Based on Transposable Elements signature in cell-free RNA"

### Early Prediction of Preeclampsia Based on Transposable Elements Deregulation in the Placenta

**Contains**

**Extended Data Fig 1–5,**

**Extended Data Table 1– 4**
Extended Data Table 1. Placental samples used for CUT&Tag and RNAseq
Extended Data Table 2: Primers used in the study
Extended Data Table 3: TE subfamilies and AUC details for final TE signatures.
Extended Data Table 4: TE signature statistics
**Source data:**

1. **Placental_RNAseq_datasets_used:** Placental RNAseq Datasets used in this study
2. **Plac_Genes_all_PEvsCT_padj<0.05**: Differentially expressed genes in PE placentas vs Control (from 5 cohorts)
3. **Plac_TE_all_PE_vs_CT_pad<0.05**: Differentially enriched TEs in PE placentas vs Control (from 5 cohorts)
4. **Plac_TE_PE+IUGRvsControl_p<0.1**: Differentially enriched TEs in PE+IUGR placentas vs Control (this study)
5. **CUT&Tag_Data_replicates:** CUT&Tag datasets generated in this study
6. **H3K27ac_bins_significant**: Differentially acetylated regions (DARs) in PE+IUGR for H3K27ac
7. **H4K16ac_bins_significant**: Differentially acetylated regions (DARs) in PE+IUGR for H4K16ac
8. **cfRNA_datsets_used**: cell-free-RNAseq Data used in this study
9. **cfRNA_TE_padj<=0.05, FC ±0.5**: Differentially enriched TEs in PE vs Control for cfRNA GA >13 weeks
10. **RRF_Var_Imps_for_133_TEs:** RRF Variable Importances for 133 differentially abundant TEs in PE-cfRNA
11. **cfRNA_TE_discovery_coh_signific**: Differentially enriched TEs in PE vs Control for cfRNA GA >13 weeks, discovery cohort
12. **cfRNA_TE_validation_coh_signifi**: Differentially enriched TEs in PE vs Control for cfRNA GA >13 weeks, validation cohort

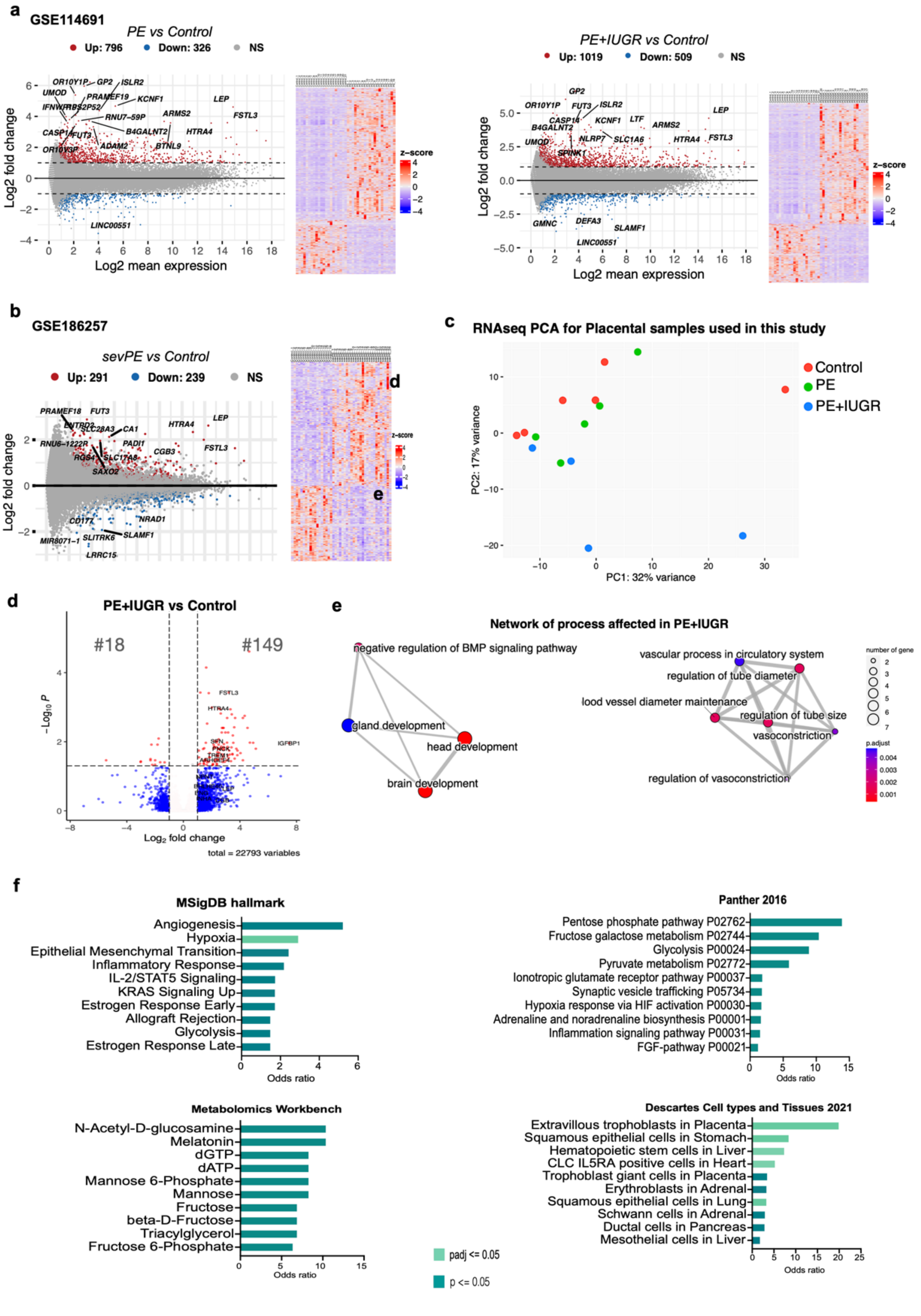

**Extended Data Fig 1: RNAseq analysis recapitulates the genes and pathways dysregulated in PE** *a) MA-plot showing DEGs (FDR or padj < 0.05, FC |1|) in PE vs Control*

*for GSE114691 (left) and PE+IUGR vs Control (right). b) Like in A for severe PE vs Control for GSE182657. Log2Mean expression on the x-axis and Log2 fold-change on the y-axis. c) Principal component analysis (PCA) plot showing the variance between samples (CT, PE and PE+IUGR) for RNA-seq. d) Volcano plot describing the differentially expressed genes (DEGs in red, FDR <0.05 and log2 foldchange ±1) in PE+IUGR (n=4) against normotensive controls (n=6). c) The network plot shows the processes significantly affected by PE+IUGR. d) Bar plots depicting odds ratio (X-axis) for significantly altered pathways and processes (Y-axis) in PE compared to Control identified using EnrichR on DEGs from fig 1c.*

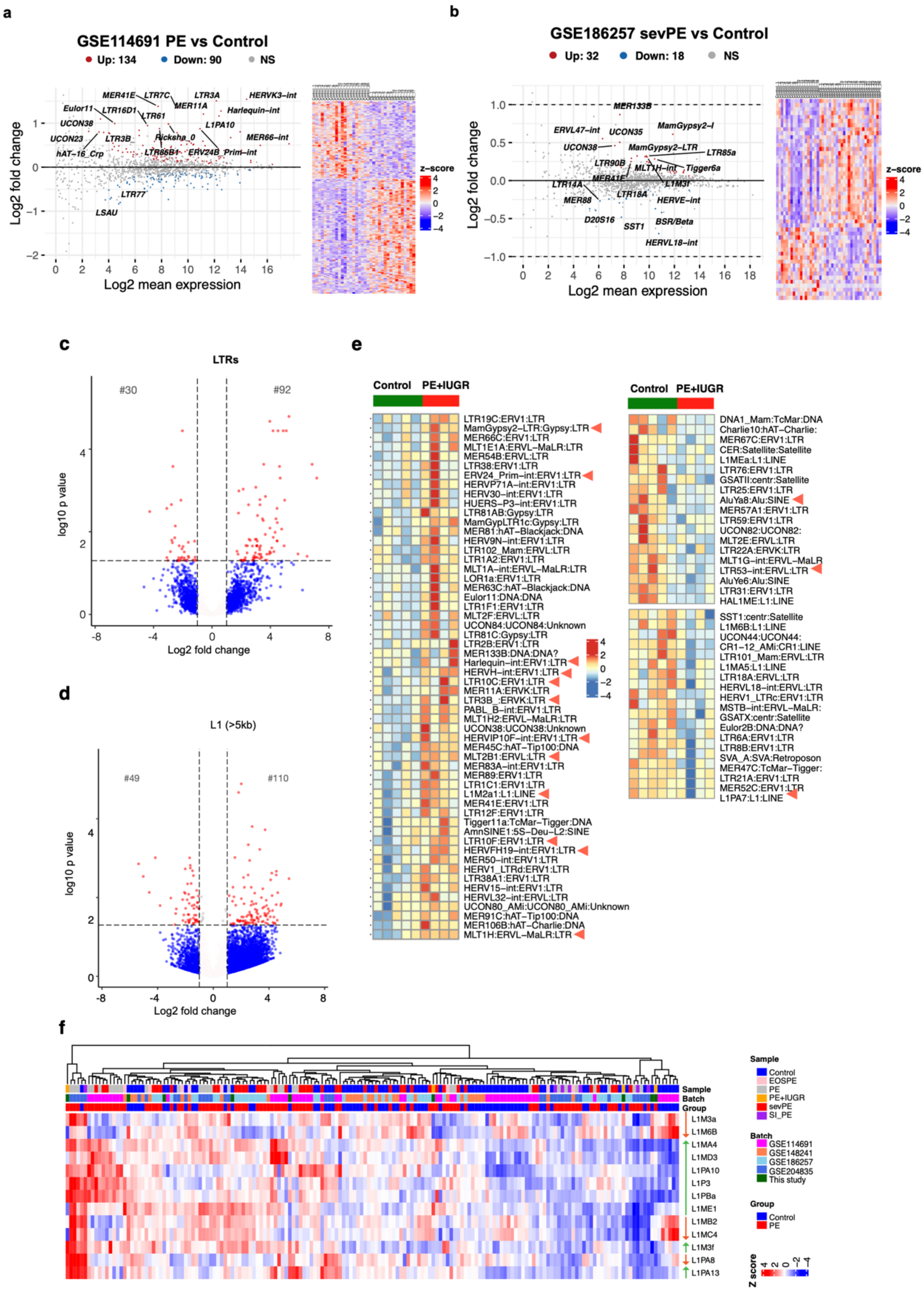

**Extended Data Fig 2: Differentially expressed TE-subfamilies in PE**

*a) MA-plot and heatmap showing differentially expressed TE-subfamilies in PE vs Control in GSE114691 ((FDR or padj < 0.05). In MA plot, Log2Mean expression on x-axis and Log2 fold-change on y-axis. b) Like in (a), for GSE186257 for TEs deregulated in severe PE vs Control. c, d) Volcano plots (padj < 0.05) showing differentially expressed LTR elements (c) and full-length L1s (d) at individual element levels between PE+IUGR and control from this study. e) Similar to Fig. 3a, but only for differentially expressed TEs in PE+IUGR (red) against control (green) heatmap vst z-score. Highlighted (triangle) TE-subfamilies show consistent expression patterns with multiple published cohorts (Fig 2a). f) Similar to 2a but only for LINE1 subfamilies that are differentially expressed with (p-value <0.05).*

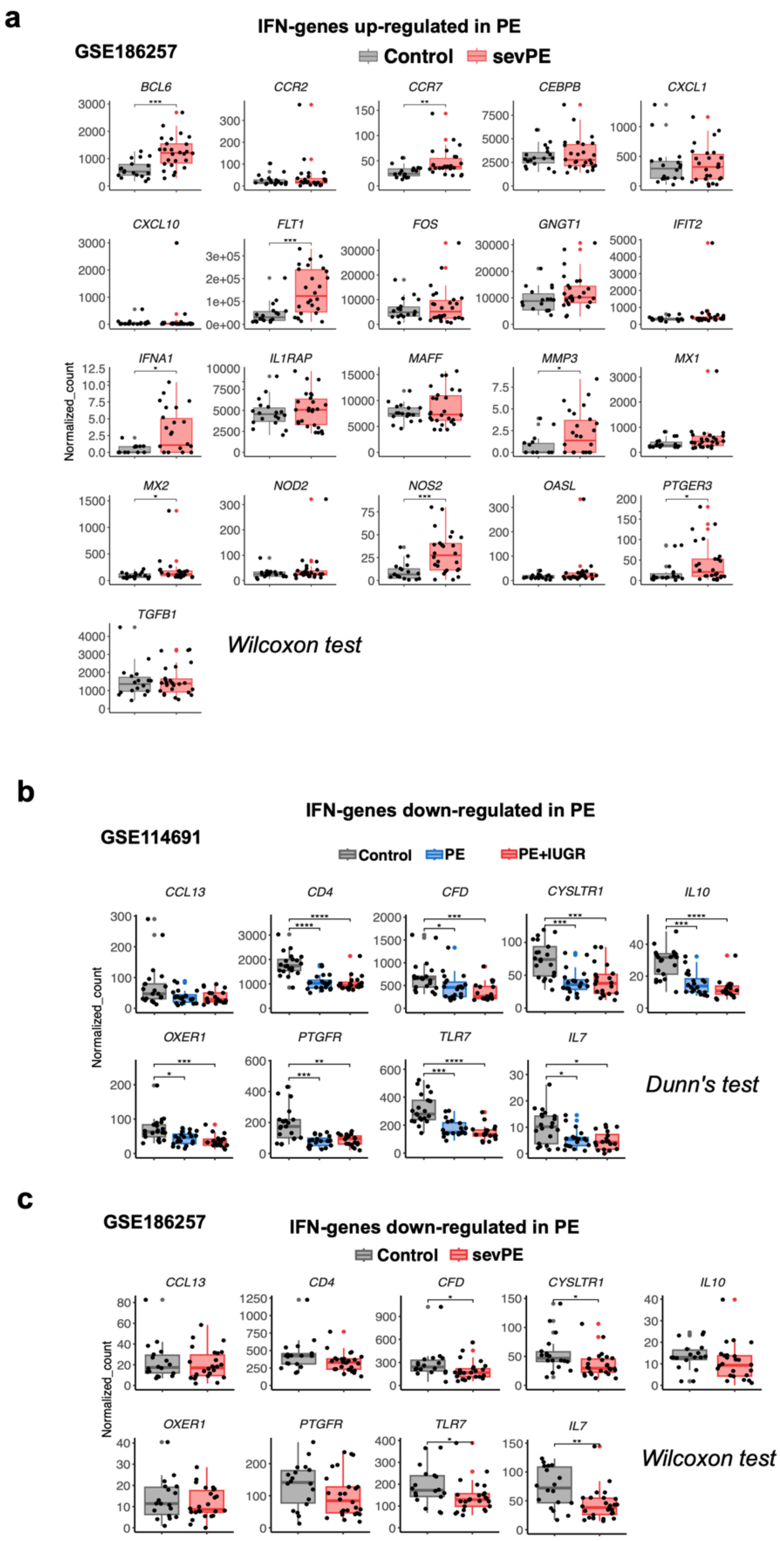

**Extended Data Fig 3: IFN-1 pathway genes are upregulated in PE**

*a) Box plots compare the expression of IFN-responsive genes up-regulated in PE (Fig. 3a) in severe PE (red) and control (grey) samples for GSE186257. b) Like in (a) for IFN-responsive genes down-regulated in PE (Fig. 3a) for GSE114691 (Control in grey, PE in blue and PE+IUGR in red). Adjusted p-values represented for significance for Dunn's test for multiple comparisons. c) Like in (b) for IFN-responsive genes down-regulated in PE (Fig. 3a) for GSE186257 (Control in grey, severe PE in red). Adjusted p-values represented for significance for Wilcoxon test.*

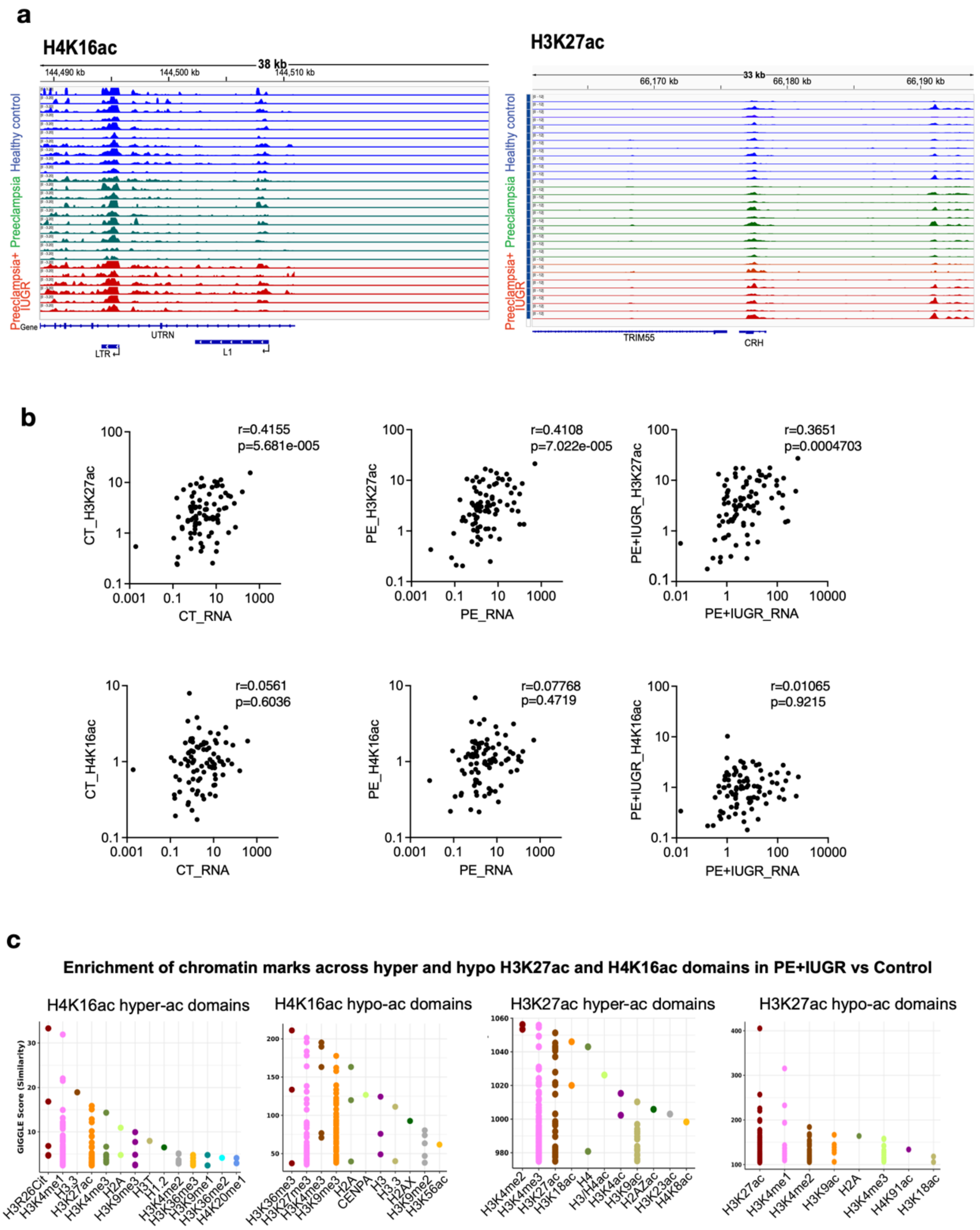

**Extended Data Fig. 4: CUT&Tag data from individual samples** *a) Genome browser tracks for H4K16ac (left) and H3K27ac (right) CUT&Tag data from healthy control, PE and PE+IUGR. Two replicates of CUT&Tag data were generated per placental sample, data*

*from individual replicates are shown. b) Correlation plots for Histone acetylation levels (y-axis, CPM) and RNA-seq (x-axis, RPKM) for genes upregulated in Preeclampsia compared for CT (left), PE (middle) and PE+IUGR (right) for H3K27ac (top panel) and H4K16ac (bottom panel).* c*) Distribution of H4K16 hyper- and hypo-acetylated domains based on the similarity profile for multiple histone modifications (x-axis) across multiple cell lines based on the GIGGLE score (detailed in methods) for similarity (y-axis) obtained with Cistrome database.*

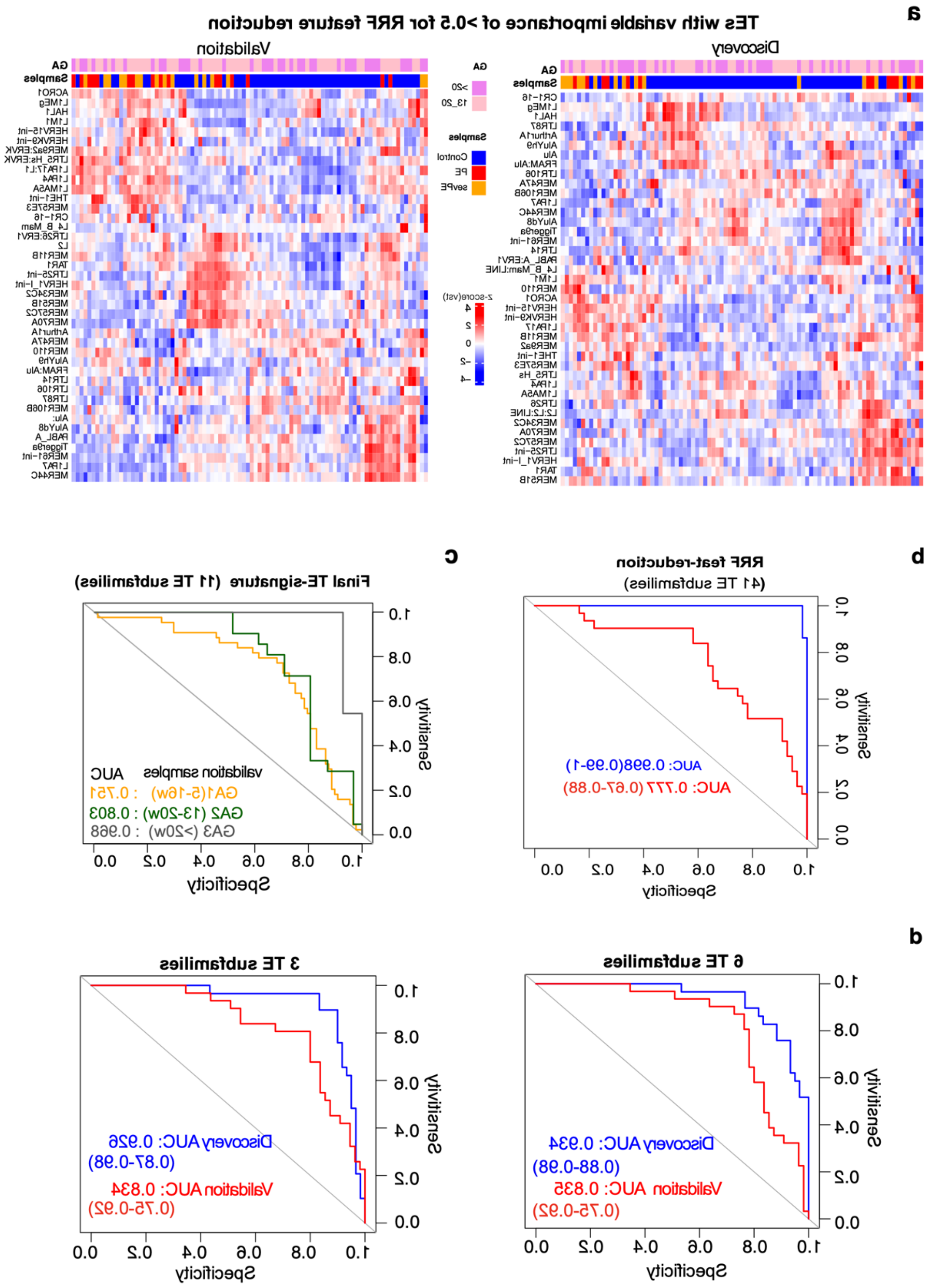

**Extended Data Fig 5: LDA modelling of TE signatures effectively predicts PE**

*a) Unsupervised clustering of heatmaps for controls, PE and severe-PE cfRNAseq (vst normalised z-scores) samples from Fig. 5a for the 41 TE subfamilies with variable importance of >0.5 (Fig. 6a) for regularised random forest (RRF) feature reduction. Discovery (n=89, left) and validation samples (n=84, right). b) The receiver operating characteristic (ROC) curve shows AUC values (with 95% CI) for TEs from (a) to the model using the LDA algorithm for discovery (blue) and validation (red) samples. c) ROC curve showing AUC values (with 95% CI) for TE signature with 11 TE subfamilies LDA model used to predict PE for Discovery samples with gestational age (GA) 5-16 weeks (orange), GA 13-20 weeks (green) and GA > 20 weeks (grey). d) ROC curve showing AUC values (with 95% CI) for TE signature-2 (n=6 TEs, left panel) and TE signature-3 (n=3 TEs, right panel) LDA models used to predict PE for discovery (blue, GA > 13 weeks) and validation (red, GA > 13 weeks) samples respectively.*

**Extended Data Table 1. Placental samples used for CUT&Tag and RNAseq**

| | **Control** | **PE** | **EOPE & PE with IUGR)** |
|---|---|---|---|
| **Number of participants** | **6** | **5** | **4** |
| Age of mothers (years) median (IQR) | 33 (30 – 34) | 32 (24 – 35) | 21 (21 - 30) |
| BMI (kg/m$^2$) median (IQR) | 27-9 (22.4-36.3) | 26.0(22.9-28.2) | 25.4 (23.5-34.8) |
| Ethnicity | 5 Asian, 1 White | 4 Asian, 1 Afro-Caribbean | 3 Asian, 1 White |
| Type of delivery | 4 EMCS – 2 vaginal | 2 EMCS – 3 vaginal | 3 EMCS – 1 vaginal |
| $MgSO_4$ (n and %) | 0 (0%) | 0 (0%) | 2/4 (50%) |
| Antenatal steroid (n and %) | 0 (0%) | 0 (0%) | 2 /4 (50%) |
| Urinary protein creatinine ratio (mg/mmol) median (IQR) | 0 (0 – 16.5) | 82.0 (42.7 – 363.5) | 505.6 (373.3 – 731.5) * |
| Another comorbidity | no | no | One had GDM |
| Gestational age (weeks) median (IQR) | 38.1 (37– 39) | 38.3 (37.9 – 39.4) | 33.4 (29.5 – 37.5) |
| Birth weight (grams) median (IQR) | 2895 (1979 - 3605) | 3020 (2795 - 3440) | 1720 (898 - 2332) |
| Need for active resuscitation(n) | 0 | 0 | 3/4 received respiratory support |
| APGAR score at 1 minute median (IQR) | 9 (9-9) | 9 (8-9) | 5 (2-8) |
| APGAR score at 5 minutes median (IQR) | 10 (10-10) | 10 (9-10) | 8 (6-10) |
| Sex (M: F) | 5 :1 | 4:1 | 3:1 |

PE, PE; IUGR, intrauterine growth restriction; EMCS, emergency caesarean section; IQR, interquartile range; GDM, gestational diabetes mellitus; * statistically significant different across study groups using nonparametric Kruskal Wallis test.

#### Extended Data Table 2. Primers used in this study

| Primer name | Forward (5'-3') | Reverse (5'-3') | Reference |
|---|---|---|---|
| GAPDH | ACCCAGAAGACTGTGGATGG | TTCTAGACGGCAGGTCAGGT | |
| PSIP/p75 | TGCTTTTCCAGACATGGTTGT | CCCACAAACAGTGAAAAGACAG | |
| HERVK9 | GGCTGGCAATAATACCTGGATG | CACCTCTGACTGTTCTGCAATG | [71] |
| IFIT1 | CCTGAAAGGCCAGAATGAGG | TCCACCTTGTCCAGGTAAGT | [72] |
| IFIT2 | ACTATGCCTGGGTCTACTATCA | TCAAGCTCTGGACTCTCAATTC | [72] |
| IFNA Common primers | TTGATGGCAACCAGTTCCAG | TCATCCCAAGCAGCAGATGA<br>TGTTCCCAAGCAGCAGATGA | [54] |
| L1 5'UTR | GCCAAGATGGCCGAATAGGA | AAATCACCCGTCTTCTGCGT | [54] |
| L1 ORF1 | ACCTGAAAGTGACGGGGAGA | CCTGCCTTGCTAGATTGGGG | [54] |

#### Extended Data Table 3. TE subfamilies and AUC details for final TE signatures.

| TE signatures | TE subfamilies | PE prediction AUC 95% CI (range) |
|---|---|---|
| TE signature | HERVK9-int, MER11B, L1PA6, MIR3, L1M7, MLT2D, MER61-int, L1PA7, AluSx4, MLT2B3, HAL1 | Discovery: 0.974(0.89–0.99)<br>Validation: 0.879 (0.81–0.95) |
| Subset of 6 TEs | HERVK9-int, MER11B, L1PA6, MIR3, L1M7, MLT2B3 | Discovery: 0.934 (0.88-0.98)<br>Validation: 0.835(0.75-0.92) |
| Subset of 3 TEs | HERVK9-int, MER11B, L1PA7 | Discovery: 0.926 (0.87-0.98)<br>Validation: 0.834(0.75-0.92) |

#### Extended Data Table 4: TE signature statistics

| Discovery | Control | PE | acc | sens | spec | PPV | NPV |
|---|---|---|---|---|---|---|---|
| Control (predicted) | 55 | 4 | 0.90 | 0.86 | 0.92 | 0.83 | 0.93 |
| PE (predicted) | 5 | 25 | | | | | |
| Validation | Control | PE | | | | | |
| Control (predicted) | 46 | 6 | 0.83 | 0.81 | 0.84 | 0.74 | 0.88 |
| PE (predicted) | 9 | 25 | | | | | |
| GA 13-20 weeks | Control | PE | | | | | |
| Control (predicted) | 23 | 6 | 0.73 | 0.71 | 0.74 | 0.65 | 0.79 |
| PE (predicted) | 8 | 15 | | | | | |
| GA >20 weeks | Control | PE | | | | | |
| Control (predicted) | 26 | 1 | 0.92 | 0.91 | 0.93 | 0.83 | 0.96 |
| PE (predicted) | 2 | 10 | | | | | |
| GA 5-16 weeks | Control | PE | | | | | |
| Control (predicted) | 67 | 16 | 0.72 | 0.64 | 0.76 | 0.57 | 0.81 |
| PE (predicted) | 21 | 28 | | | | | |
